# Genome-scale functional mapping of the mammalian whole brain with in vivo Perturb-seq

**DOI:** 10.64898/2026.03.16.711480

**Authors:** Tuo Shi, Maria Korshunova, Seoyeon Kim, David DeTomaso, Xinhe Zheng, Lavanya Vishvanath, Thokozile Nyasulu, Nhan Huynh, Alexander Sun, Patrick C. Thompson, Yifan Zhang, Emilie M. Wigdor, Narjes Rohani, Salma Ali, Huixian Qiu, Michael Geralt, Ziyan Zhao, Sara Rabhi, Zizhen Yao, Cindy TJ van Velthoven, Joseph R. Nery, Rosa Gomez Castanon, Severin Dicks, Tiffany J. Chen, Joseph R. Ecker, Hongkui Zeng, Grace XY Zheng, Stephan J. Sanders, Laksshman Sundaram, Xin Jin

## Abstract

Functional genomics studies have provided critical insights into cell type-specific gene regulatory programs, but to date most have been conducted in wild-type tissues or cell cultures. Here, we present a gene expression functional atlas across the mouse brain. We use an enhanced in vivo Perturb-seq platform to analyze transcriptome-wide responses to loss of 1,947 disease-associated genes, profiling over 7.7 million cells spanning major brain regions and neuronal populations. We find striking cell-type-specific essentiality and transcriptional programs and show that closely related disease genes such as two NMDA receptor subunits can drive opposing transcriptional programs. Together, this work reveals insights into the genetics and mechanisms of neurodevelopmental, psychiatric, and neurodegenerative diseases in vivo, paving the way for the design of future genetic medicine.

## Main text

Probing gene function often begins with understanding the consequences of loss of the gene in question. shRNA and CRISPR knockout screens have scaled this principle to interrogate the entire genome and have enabled high-throughput functional screening in mammalian cells, greatly advancing our ability to study gene function systematically^1^. These screens, however, are largely limited to cell culture, precluding an understanding of how cell type-specific context affects gene function. This is particularly challenging for studying how mutations give rise to neurodevelopmental and psychiatric diseases, where the myriad cell types of the brain and their connections create a complex genetic landscape that cannot be fully recapitulated in vitro^2-5^. Here, we generated a genome-scale, in vivo perturbation atlas targeting 1,947 disease-relevant genes across the whole mouse brain, profiling their cell-autonomous phenotypes with snRNA-seq from 7.7 million cells in postnatal brains. Genetic perturbations exhibit both shared and cell-type-restricted effects across the brain transcriptome, with several key transcriptional responses concentrated in specific neuronal subclasses. Distinct sets of human disease genes resolve preferentially to different brain regions and cell types, revealing prominent dysregulation beyond the cortex, including midbrain and thalamic glutamatergic neuronal types. This atlas links gene function to cell type-specific outcomes in the mammalian brain, establishing a framework for mechanistically connecting disease-associated genetic risk to the cellular programs and neuronal contexts in which it acts.

### Scaling in vivo Perturb-seq to whole-brain coverage with expanded throughput

To overcome the scalability constraints of in vivo Perturb-seq, we developed a high-throughput whole-brain platform that integrates CRISPR perturbations with single-nucleus RNA sequencing (snRNA-seq) (Fig. 1A-B). We injected 74 young (P16) Cas9 transgenic mice with a pooled AAV library encoding CRISPR guide RNAs (gRNAs) targeting 1,947 genes and non- and safe-targeting control gRNAs (Table S1). The 1,947 target genes were selected based on their broad neuronal expression in both mouse and human brain (Fig. 1C, S1A-B) and strong relevance to human disease (see Methods), with prioritization of transcription factors and receptors given their central roles in gene regulation^2,6^. To ensure the gene editing efficacy, we tested and validated gRNA activities of 15 target genes in HT22-Cas9 cells, which had an average editing efficiency of 46.5% (Fig. 1D, S1C-D; Table S2).

**Figure 1.**
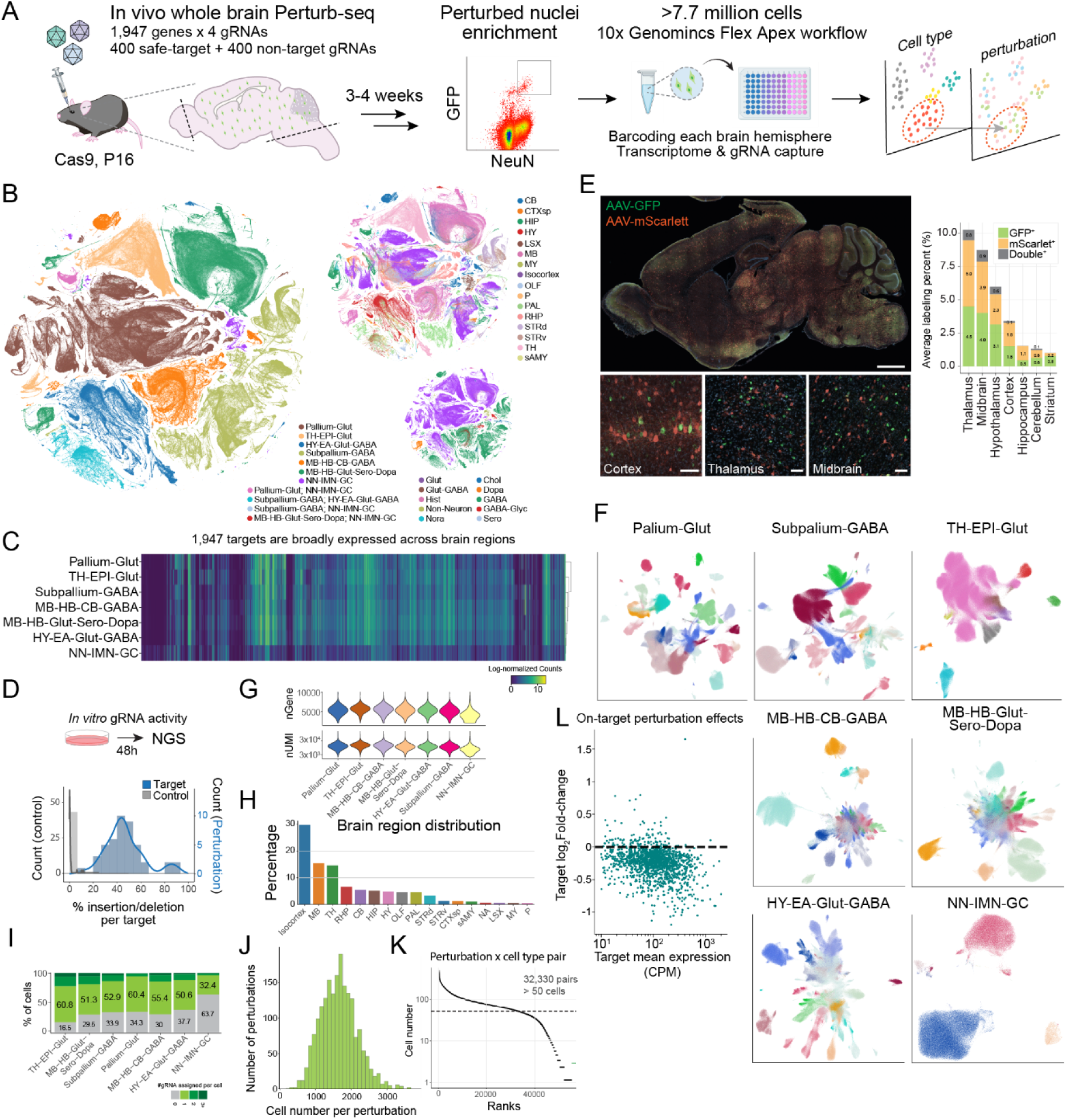
Genome-scale in vivo Perturb-seq achieves whole-brain coverage with high gRNA fidelity and cell type resolution. (**A**) Schematic of mouse whole brain in vivo Perturb-seq using 10x Genomics Flex Apex platform. (**B**) UMAP of whole brain in vivo Perturb-seq dataset encompassing 7.7 million sequenced nuclei, colored by developmental neighborhoods, anatomical region, and neurotransmitter type, inferred using MapMyCells^86^. (**C**) Heatmap of the gene expression levels of 1,947 neurodevelopmental disease-associated risk genes in non-targeting control nuclei across different developmental neighborhoods. (**D**) Histogram of in vitro gRNA activity distribution of 45 selected gRNAs (15 genes, 3 gRNAs per gene) compared to safe-targeting controls by insertion-deletion analysis. (**E**) Immunofluorescence image of sagittal section of a P37 mouse brain retro-orbitally administered with 6e8 total vg per gram of body weight of AAV PHP.eB encoding either GFP or mScarlet (1:1 ratio) (scale bar, 1 mm), accompanied by zoomed in images to show representative MOI in each major brain region (scale bar, 50 μm), and stacked bar plot quantifying GFP and mScarlet viral labeling efficiency as well as double labeling rate. (**F**) UMAPs of whole brain in vivo Perturb-seq dataset separated by neighborhoods, colored by inferred cell subclass using MapMyCells^86^. (**G**) Violin plots of number of genes and RNA UMIs recovered per nucleus from each developmental neighborhood. (**H**) Ranked bar plot showing proportion of sampled nuclei by brain region. (**I**) Stacked bar plot showing percentage of nuclei with no guide, single, double, or multiple guide assignment within each developmental neighborhood. (**J**) Histogram of total nuclei number distribution of nuclei recovered per perturbation. (**K**) Ranked dot plot of nuclei number in each perturbation and cell type pair and the minimum cell number cut off for perturbation and cell type pair for downstream analyses (dashed line). (**L**) Scatter plot of weighted mean log fold-changes of target genes across all cell types against their weighted mean expression levels in non-targeting control nuclei.

We used a previously established transposon system to stabilize vector expression, as well as an enhanced capture strategy to enrich for gRNA recovery via nuclear anchoring of the transcript^7-9^. Vector dosage was titrated to maximize nuclei labeling while minimizing multiple transduction events across the entire mouse brain (Fig. 1E, S2A-B). Three to four weeks later, we harvested whole brains, removed olfactory bulbs and hindbrains, and extracted nuclei from each hemisphere for FACS enrichment. Across the 74 brains, on average, 1.6% GFP^+^/NeuN^+^ nuclei were enriched from sparsely labeled brains (Fig. S2C-E; Table S3). We prepared snRNA-seq libraries from these nuclei using the 10x Genomics Flex Apex platform, enabling cell- and sample-level barcoding and banking of millions of perturbed nuclei, while preserving each cell’s animal identity.

We recovered 7,720,300 high-quality NeuN+ nuclei corresponding to the reference neuronal cell types and brain regions^6^ (Fig. 1B, S2F). After quality control, across 34 classes and 331 subclasses, these cells can be further organized into lineage- and anatomy-informed neighborhoods reflecting their regional identities: Pallium-Glut, containing glutamatergic neurons across all pallium structures (including the isocortex, HPF, OLF and CTXsp); Subpallium-GABA, containing subpallium-derived GABAergic neurons across cortical and striatal regions; HY-EA-Glut-GABA, encompassing glutamatergic and GABAergic neurons of the hypothalamus and extended amygdala; TH-EPI-Glut, containing all thalamic and epithalamic glutamatergic neurons; MB-HB-Glut-Sero-Dopa, containing midbrain and hindbrain glutamatergic, serotonergic and dopaminergic neurons; MB-HB-CB-GABA, containing all GABAergic neurons from midbrain, hindbrain and cerebellum; and NN-IMN-GC, containing distinct non-neuronal cell types, immature neurons, and granule cell types (Fig. 1F, S3A). We performed an independent dissection experiment to validate that anatomically defined brain regions and their constituent cell types can be accurately recovered using this mapping strategy (Fig. S1E-H). As cell type abundance varies substantially across regions, with some areas, such as the hypothalamus, exhibiting high diversity and granularity, we consolidated related subclasses into 29 binned neuronal and 2 binned non-neuronal groups (Table S4). Overall, nuclei retained a median of 12,146-15,548 RNA UMI/nucleus and enriched in neuronal representation of all the major brain regions (Fig. 1G-H, S3B-E).

From the 7.7 million nuclei, we achieved robust gRNA detection and confidently assigned perturbation identities to over 66.49% of the nuclei in most cell types, with 50% of total nuclei carrying a single perturbation (Fig. 1I, S4). gRNA capture efficiency is consistent across cell types, batches, and animals, with reduced labeling primarily in hippocampus, cerebellum, and glial populations, consistent with AAV tropism and NeuN enrichment sorting biases. This strategy yielded a median of 1,795 nuclei per perturbation and over 32,330 pairs of perturbation-by-cell type containing over 50 nuclei, enabling cell-type-resolved perturbation analyses across the whole brain (Fig. 1J-K, S4C). On average, in vivo CRISPR perturbation led to 18.1% downregulation per gRNA for targets with an expression of at least 100 CPM (Fig. 1L). As frameshift allele transcripts could escape complete nonsense-mediated decay, mutant transcripts may be readout by the FLEX snRNAseq despite loss of functional protein, indicating that this is likely an underestimate of the knockout^10,11^. Together, these results establish a high-quality, genome-scale in vivo perturbation resource spanning the entire mouse brain.

### Context-dependent genetic essentiality in neuronal types

Genetic perturbation can alter cell fate specification or survival, ultimately shifting the proportional representation of cell types within a tissue^12,13^. To systematically interrogate how loss of function of each of the 1,947 targeted genes affects the representation of the 29 binned neuronal types in our analysis, we compared the gRNA composition within each cell type to the expected distribution based on the initial AAV pool. Because the neuronal populations assayed are postmitotic, perturbations that confer a growth advantage cannot expand a given type; accordingly, we observed only depletion – never enrichment – of specific perturbation-cell type pairs. We applied Fisher’s exact tests to identify perturbation–cell type pairs exhibiting significant dropout, with MAGeCK-based analysis yielding concordant results^14^ (Fig. S5; Table S5).

Across all perturbation-cell type combinations, a subset of genes drove significant depletion in one or more neuronal types (Fig. 2A). Notably, the depleted genes were not uniformly shared across cell types: certain perturbations, such as those targeting the RNA helicase *Ddx39b* and molecular chaperones *Hspa5* (or BiP) and *Hspa8* (or Hsc70), caused broad depletion across many neuronal classes, consistent with their known roles in fundamental cellular processes including mRNA export, proteostasis, and protein quality control. In contrast, other perturbations produced cell-type-restricted depletion; for example, genes involved in vesicular acidification (*Atp6v1b2*, *Atp6v1a*) and lipid transport (*Pafah1b1*) preferentially depleted specific populations, suggesting that these pathways are differentially required for the survival or maintenance of distinct neuronal identities. Among the cell types surveyed, non-GABAergic midbrain neurons (MB-HB-Glut-Sero-Dopa) and glutamatergic thalamic neurons (TH-EPI-Glut) appeared most susceptible to perturbation-induced dropout, whereas others (Pallium-Glut, Subpallium-GABA, and HY-EA-Glut-GABA) were comparatively resilient. To confirm that observed depletions reflect genuine biological effects rather than single-guide artifacts, we examined gRNA-level consistency for each significant perturbation-cell type pair: the majority were supported by two or more independently depleted gRNAs (Fig. 2B), and individual gRNA-level effect sizes were concordant within each pair (Fig. 2C). Together, these results reveal a landscape of context-dependent genetic essentiality in which both broadly required housekeeping pathways and cell-type-specific programs govern neuronal survival in vivo.

**Figure 2.**
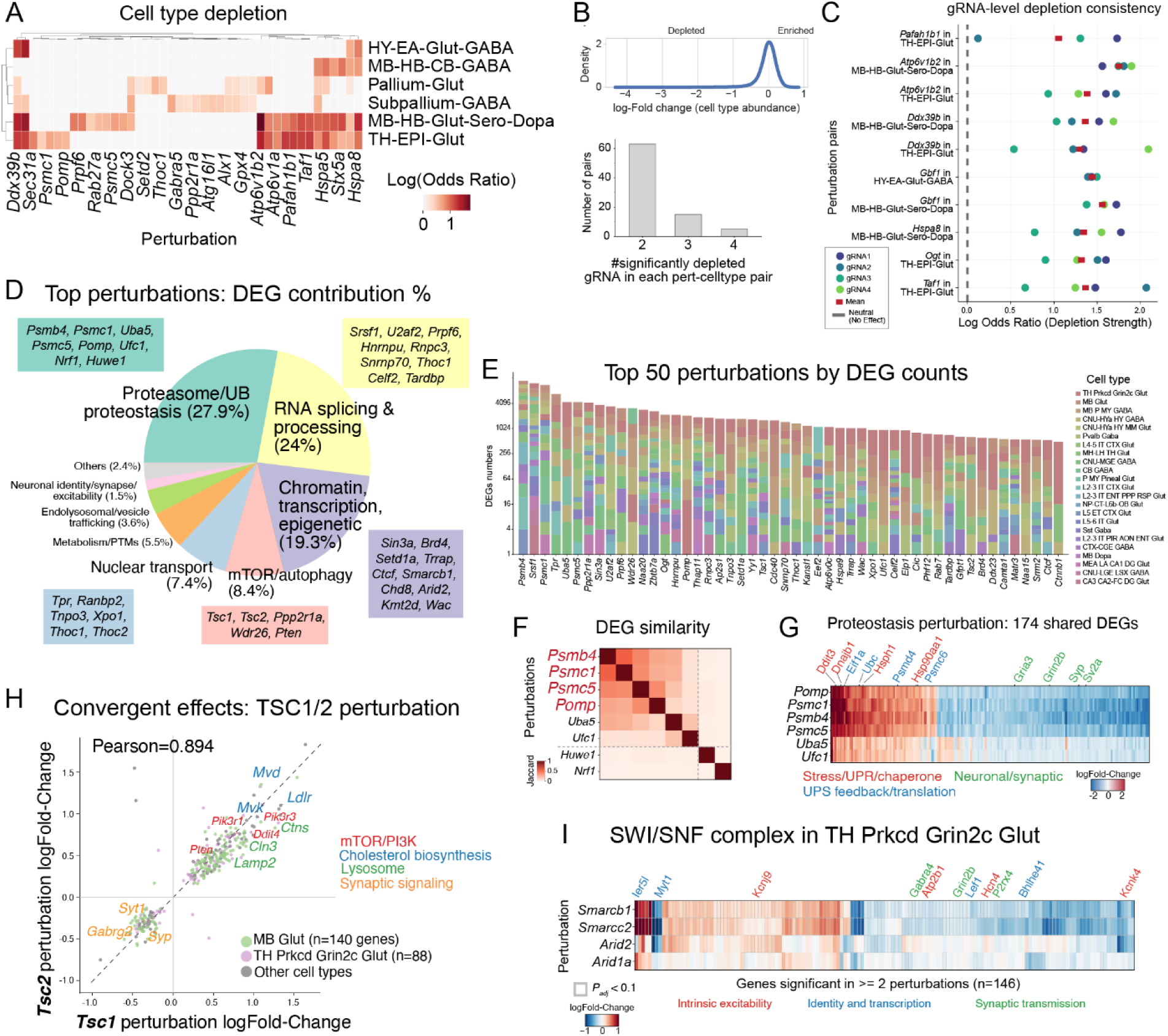
Context-dependent genetic essentiality and transcriptional consequences of perturbation across neuronal types. (**A**) Heatmap of cell type depletion showing log odds ratio for significantly depleted perturbation–cell type pairs across six binned neuronal classes. (**B**) Top: density plot of log₂(cell fraction/library fraction) across all non-control perturbation-cell type pairs. (**C**) gRNA-level depletion consistency for selected perturbation–cell type pairs, showing individual gRNA log odds ratios (colored dots) and mean (red square) relative to neutral expectation (dashed line). (**D**) Pathway composition of DEGs among the top 422 perturbations, with representative genes listed for each category. (**E**) Stacked bar plot of DEG counts across cell types for the top 50 perturbations ranked by total DEG number. (**F**) Jaccard similarity of DEG sets among proteostasis-related perturbations, showing high overlap among proteasome and ubiquitin pathway components. (**G**) Heatmap of log fold-changes for 174 shared DEGs across proteostasis perturbations, organized by functional category (stress/UPR/chaperone, UPS feedback/translation, neuronal/synaptic). (**H**) Scatter plot of DEG log fold-changes for *Tsc1* versus *Tsc2* perturbations (Pearson R = 0.894), with shared DEGs colored by pathway (mTOR/PI3K, cholesterol biosynthesis, lysosome, synaptic signaling). Cell-type-specific effects highlighted for MB Glut and TH Prkcd Grin2c Glut. (**I**) Heatmap of log fold-changes for SWI/SNF complex perturbations (*Smarcb1*, *Smarcc2*, *Arid2*, *Arid1a*) in TH Prkcd Grin2c Glut neurons, showing DEGs significant in two or more perturbations (n = 146), organized by functional category (intrinsic excitability, identity and transcription, synaptic transmission).

### Core proteostasis and other regulatory pathways drive widespread neuronal perturbation signatures

We next examined the global transcriptional consequences of perturbation across cell types. We computed differentially expressed genes (DEGs) for each of the 1,947 perturbations within each of the 31 binned cell types relative to non-targeting controls using a Wilcoxon rank-sum test (see Methods). Significance was determined after correction for multiple testing across genes using a Benjamini-Hochberg procedure^15^, providing a conservative threshold given the large number of perturbation-cell type comparisons (Table S6). We found that the perturbations with the highest total number of DEGs were most enriched for genes involved in protein homeostasis and degradation (27.9% of the DEGs), RNA splicing and processing (24%), chromatin and transcriptional regulation (19.3%), and mTOR/autophagy (8.4%) (Fig. 2D); these DEGs span all 23 cell types (Fig. 2E). These pathway categories were significantly overrepresented relative to their frequency among all perturbed genes (Fig. S6A), indicating that specific molecular systems disproportionately cause wide-spread transcriptional changes in neurons.

We next examined both the potency (number of DEGs) and magnitude (mean absolute log-Fold-change) of each perturbation across cell types (Fig. S6B-C; Table S6). Individual perturbations did not elicit uniform responses across cell types; some cell types consistently showed stronger responses across many perturbations, whereas others were comparatively less responsive. Perturbations targeting genes involved in RNA processing, transcription, proteostasis, and vesicle trafficking tended to produce larger numbers of DEGs overall, though not in all the cell types. Across perturbations, the magnitude of expression shifts was typically modest and only weakly related to potency (Fig. S6C). Together, these results indicate that the transcriptomic consequences of gene loss-of-function in postmitotic neurons are shaped both by the functional role of the perturbed gene and by the identity of the cell type in which it is perturbed.

We also observed that perturbations of known complexes converged on shared DEG programs. Similarity analysis of DEG sets revealed clusters of genes with overlapping transcriptional consequences (Fig. 2F-I). Notably, perturbation of proteasome components (*Psmc1*, *Psmb4*, *Psmc5*, *Pomp*) and UFMylation pathway genes (*Uba5*, *Ufc1*) induced highly concordant responses, sharing 174 DEGs across all six perturbations (Fig. 2F-G). These shared signatures were characterized by upregulation of unfolded protein response and stress-related genes and coordinated downregulation of neuronal and synaptic programs, consistent with disrupted proteostasis impairing neuronal function. We also observed convergent transcriptional effects among paralogs such as *Tsc1* and *Tsc2*, whose perturbations showed highly correlated effect sizes across cell types and enriched downstream mTOR/PI3K, cholesterol biosynthesis, lysosomal pathways, and downregulation of synaptic genes (Fig. 2H). Additionally, perturbation of core SWI/SNF scaffold subunits (*Smarcb1* and *Smarcc2*) in the thalamic glutamatergic neurons produced concordant transcriptional changes: downregulation of synaptic transmission and intrinsic excitability components and upregulation of developmental and state-associated transcriptional regulators (Fig. 2I). These patterns suggest that core SWI/SNF activity contributes to maintaining neuronal functional gene programs in vivo, with partial but weaker effects observed for perturbations of *Arid2* and *Arid1a*.

Lastly, we validated our DEGs against existing published germline knockout or conditional knockout mouse data harboring the same perturbations (Fig. S7). Across the board, we observed concordant gene expression changes in *Chd8* (Pearson R = 0.85)^16^, *Mef2c* (Pearson R = 0.7)^17^, *Ctcf* (Pearson R = 0.48)^18^, *Grin2a* (Pearson R = 0.63)^19^, and *Myt1l* (Pearson R = 0.81)^20^. Notably, our neuronal *Mef2c* perturbation did not correlate with knockout data from iPSC-derived microglia (Fig. S7B, Pearson R = 0.05)^21^, emphasizing the importance of cellular context in interpreting perturbation consequences. Together, at the genome scale and preserving cellular context in vivo, we resolve these convergent vulnerabilities at single-cell resolution, revealing signatures consistent with known biology.

### Perturbation landscape reveals modular organization of transcriptional responses to KOs

To determine whether perturbations with strong transcriptomic effects cluster together, we computed pairwise energy distances between all effective perturbations (Fig. 3A, S8; Table S7; see Methods). Rather than a continuum of independent effects, the perturbation landscape was modular: perturbations with high DEG counts clustered together, indicating that transcriptionally potent perturbations shift cells into similar transcriptomic states. For example, chromatin and transcriptional regulators (*Kmt2a*, *Arid2*, *Chd2*, *Smarcb1*) clustered together, as did RNA-processing factors (*Snrnp70*, *Srsf1*, *Hnrnpu*, *Tardbp*) and genes governing neuronal excitability and synaptic signaling (*Grin2a*, *Grin2b*, *Kcnb1*).

**Figure 3.**
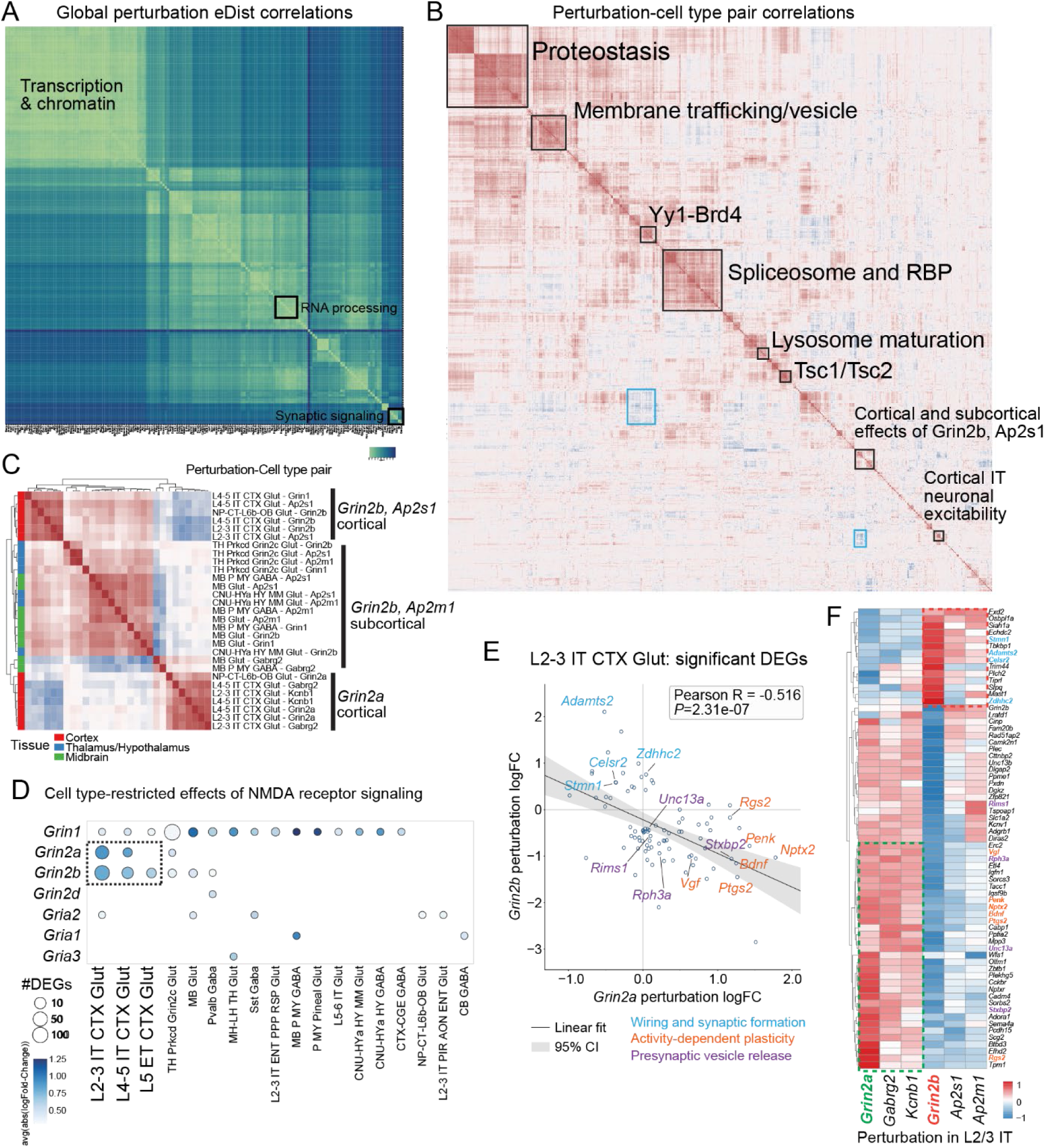
Perturbation landscape reveals modular organization of transcriptional responses. (**A**) Pairwise energy distance matrix across all effective perturbations, clustered hierarchically, showing modular structure among transcriptionally potent perturbations. (**B**) Pairwise cosine similarity of signed DEG vectors across perturbations, with hierarchical clustering revealing biologically coherent modules. (**C**) Sub-clustering of the neuronal excitability and synaptic signaling module, resolving *Grin2a*- and *Grin2b*-associated perturbation groups into distinct clusters. (**D**) Cell-type-specific examples of *Grin2a* and *Grin2b* perturbation effects (DEGs padj < 0.1), illustrating divergent transcriptional programs across neuronal populations. (**E**) Scatter plot of shared DEG (padj < 0.1) effect sizes between *Grin2a* and *Grin2b* perturbations in L2/3 IT cortical glutamatergic neurons, showing anti-correlated transcriptional responses (Pearson R = −0.516). (**F**) Heatmap of selected DEGs (padj < 0.05) showing opposing regulation between *Grin2a* and *Grin2b* perturbations, including wiring and synaptic formation genes, activity-dependent plasticity genes, and presynaptic vesicle release machinery genes.

Beyond the global clustering, we found that within each brain region and neighborhood, the clustering patterns are also distinct (Fig. S8; Table S7), which motivated us to investigate the relationship across the cell-type-perturbation pairs. To resolve finer architecture, we used a variational autoencoder to model perturbations across the full dataset as summarized effect vectors and performed hierarchical clustering of 1,218 perturbation-cell-type pairs, revealing tightly organized modules with distinct biological signatures (Fig. 3B, S9; Table S7; see Methods). The most internally concordant module comprised perturbations involved in proteostasis and mitochondrial homeostasis, including molecular chaperones (*Psmb4*, *Pomp*) (Fig. S9A) and mitochondrial quality-control regulators (*Hspa5*, *Hspa8*, *Hspa9*, *Lonp1*, *Opa1*, *Dnm1l*) (Fig. S9B), which induced nearly identical transcriptional responses across diverse cell types. A second tightly correlated module encompassed membrane trafficking and vesicle homeostasis genes, including endosomal and secretory machinery components such as *Vps37a*, *Vps53*, *Snf8*, *Atp6v0c*, *Atp6v1a*, *Sec31a*, *Pitpna*, and *Tbcd* (Fig. 3B, S9C): disruption of vesicle trafficking, endosomal maturation, and V-ATPase-mediated acidification thus converges onto a shared transcriptional program.

Beyond internal module concordance, we also observed structured antagonistic relationships between modules. Although core spliceosome and RNA-binding proteins – including *Prpf6*, *Snrnp70*, *Rnpc3*, *U2af2*, and *Srrm2* – formed another tight module, consistent with global disruption of mRNA splicing and RNA metabolism^22-25^ (Fig. 3B, S9D), their perturbation effects formed antagonistic relationships with vesicle trafficking and homeostasis modules (Fig. 3B, S9E). Chromatin remodeling and transcriptional regulators are organized into several related but distinct sub-modules. One included SWI/SNF components and histone modifiers such as *Kmt2a*, *Kmt2e*, *Huwe1*, *Arid2*, *Cic*, and *Tbl1xr1*^26-30^ (Fig. S9F); another encompassed regulators associated with enhancer activity and transcriptional elongation, including *Yy1* and *Brd4*^31,32^ (Fig. 3B, S9G). Perturbations within these chromatin modules produced related transcriptional responses but with greater context dependence compared to the proteostasis module, and clustering revealed an antagonistic axis between *Yy1*/*Brd4*-associated transcriptional activation and repressive chromatin regulators, indicating that perturbation of transcriptional control can drive cells toward opposing regulatory states (Fig. S9H). These opposing relationships indicate that perturbations do not simply scale a common transcriptional response but reposition cells along structured biological axes, demonstrating that large-scale in vivo perturbation profiling can resolve the functional architecture of the genome directly in intact neural circuits. Finally, a distinct module corresponded to neuronal excitability and synaptic signaling, including *Grin2a*, *Kcnb1*, *Gabrg2*, *Snap25*, *Celf2*, and *Ppp2r1a* – ion channels, neurotransmitter receptors, vesicle fusion machinery, and signaling regulators whose perturbation induced correlated transcriptional changes consistent with disruption of activity-dependent gene expression programs^33-37^ (Fig. S9I).

### Cell type-specific glutamatergic signaling in cortical and subcortical neurons

Context-restricted perturbations were further segregated according to shared cell-type sensitivity (Fig. 3C-F). A particularly informative case involved the NMDA receptor subunits *Grin2a* and *Grin2b*, which encode closely related glutamate receptor components with partially overlapping expression patterns but distinct developmental trajectories and disease associations^38,39^. *GRIN2A* variants are primarily linked to schizophrenia^40,41^, whereas *GRIN2B* variants are strongly associated with early-onset disorders including autism spectrum disorder (ASD) and neurodevelopmental disorder (NDD)^42^. A longstanding view attributes this divergence largely to their developmental expression dynamics: *Grin2b* is highly expressed during early cortical development, whereas *Grin2a* becomes predominant postnatally as excitatory circuits mature^43,44^.

We found perturbations of these two genes produced significant transcriptional changes in overlapping neuronal populations (Fig. 3B), yet resolved into distinct clusters with different biological correlate (Fig. 3C). *Grin2a* perturbations clustered with cortical excitability-associated genes including *Kcnb1* and *Gabrg2*, forming a module enriched in cortical glutamatergic neurons: consistent with a shared role in regulating intrinsic excitability and synaptic transmission in cortical circuits^35-37^. In contrast, *Grin2b* perturbations clustered most closely with components of the AP2 adaptor complex for clathrin-mediated endocytosis (*Ap2m1*, *Ap2s1*), suggesting coupling of *Grin2b*-dependent signaling to endocytic and trafficking machinery^45,46^. Correspondingly, *Grin2b* effects were observed across both cortical and subcortical populations but with region-dependent correlation structure (Fig. 3D), implying their context-specific functional engagement.

Strikingly, transcriptional responses to *Grin2a* and *Grin2b* perturbations were frequently directionally divergent within the same cell type. In L2/3 IT cortical glutamatergic neurons, effect sizes of shared DEGs were strongly anti-correlated (Pearson R = - 0.516; Fig. 3D-E), indicating that perturbation of these closely related subunits drives opposing transcriptional programs. Several genes associated with synaptic plasticity and neuronal activity – including *Bdnf*, *Nptx2*, *Rgs2*, and *Ptgs2* – exhibited opposite regulation between the two perturbations (Fig. 3E; Fig. S10). Relative to *Grin2b* loss, *Grin2a* loss was associated with upregulation of activity-dependent genes (*Bdnf*, *Nptx2*, *Vgf*, *Penk*, *Rgs2*, *Ptgs2*) and presynaptic release machinery (*Rims1*, *Unc13a*, *Rph3a*, *Stxbp2*), supporting that *Grin2a* loss preferentially engages an activity-linked synaptic plasticity program^33,34^. *Grin2b* loss instead upregulated genes more consistent with structural remodeling and membrane regulation (*Stmn1*, *Celsr2*, *Adamts2*, *Zdhhc2*), while the activity-dependent and presynaptic genes were relatively reduced^47,48^ (Fig. 3F).

These findings suggest that, although *Grin2a* and *Grin2b* encode closely related glutamate receptor subunits, their perturbation engages distinct intracellular signaling and transcriptional programs across neuronal contexts. Thus, beyond their developmental expression differences, these two genes exert fundamentally different downstream molecular effects, providing a potential mechanistic basis for the divergent disease associations of their loss of function that cannot be learned with only wildtype data. Together, these results demonstrate that large-scale in vivo Perturb-seq resolves both shared and context-dependent transcriptional consequences of genetic perturbations across the brain, linking disease-associated genes to the specific cellular programs and neuronal contexts in which they act.

### Human disease genes converge on cell-type-specific neuronal programs

Of the 1,947 genes targeted in this screen, nearly all correspond to mouse homologs of human disease-associated genes, including 378 linked to NDD, 125 to ASD, 228 to developmental disorder (DD), and smaller subsets associated with psychiatric and neurodegenerative conditions^40,49-55^ (Fig. 4A; Table S8 and S9). In addition to disorder annotations, genes were further characterized by their tolerance to loss-of-function (LoF) variation using the LOEUF metric^56^. Gene essentiality was defined based on genomic-scale CRISPR knockout screens in human cell lines, with genes whose disruption impaired cellular viability classified as essential^57^.

**Figure 4.**
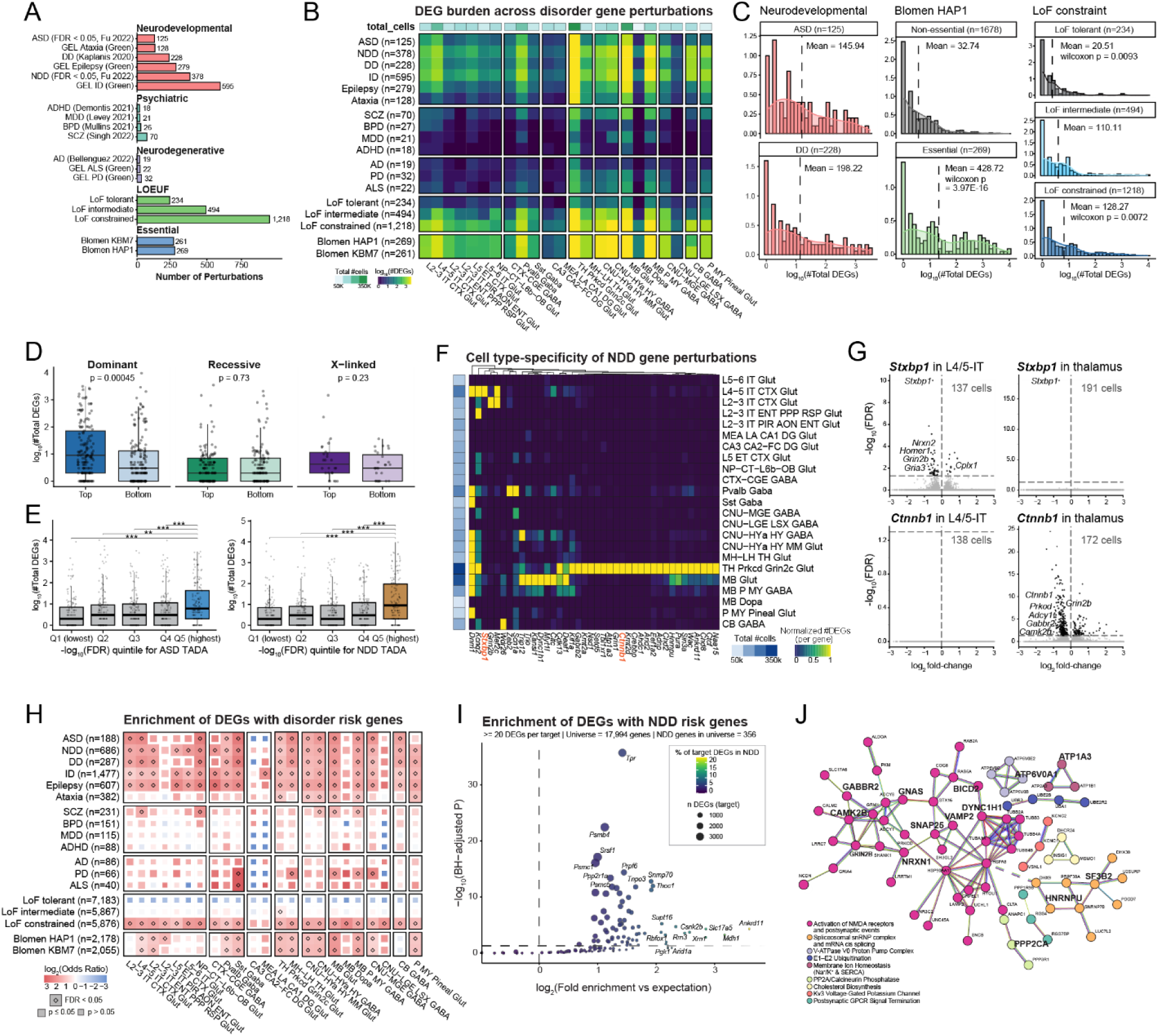
Disease-associated genes produce cell-type-specific transcriptional effects in vivo. (**A**) Number of genetic perturbations overlapping with neurodevelopmental, psychiatric, and neurodegenerative disease gene sets. (**B**) Heatmap showing DEG burden (log_10_ total DEGs, FDR < 0.05) across neuronal cell types following perturbation of disorder-associated genes, LoF constraint categories, and essential genes^57^. (**C**) Distribution of total DEG counts (log_10_ scale) for perturbations of disorder-related and functional gene categories. For Blomen HAP1 groups, differences between essential and non-essential genes were assessed using two-sided Wilcoxon rank sum tests. For LoF constraint groups, overall differences were evaluated using the Kruskal-Wallis rank sum test (p = 4.06E-06), followed by pairwise Wilcoxon rank-sum tests comparing against LoF-intermediate groups. (**D**) Total DEG counts stratified by inheritance pattern among NDD genes. Genes were divided into top and bottom halves based on dominant (dom_score > 20) and recessive (rec_score > 10) scores. X-linked scores were evaluated for genes on the X chromosome. Differences were assessed using two-sided t-tests. (**E**) Total DEG counts stratified by human genetic evidence strength (FDR quantile) for ASD/NDD genes. Genes were grouped into five quantiles based on -log10 (FDR), and differences were evaluated using two-sided t-tests (**: p-value ≤ 0.01; ***: p-value ≤ 0.001). (**F**) Heatmap showing DEG burden (FDR < 0.1) under NDD gene perturbations with the largest transcriptional effects. DEG counts were normalized per gene using min–max scaling to a 0–1 range across cell types. (**G**) Example cell-type-specific volcano plots for selected NDD gene perturbations. (**H**) Gene set enrichment test of DEGs per cell type (FDR < 0.05, absolute Log_2_ Fold Change > 0.5) with a full list of disorder risk genes, LoF constraint categories, and essential genes^57^. One-sided Fisher’s exact test was used to compute statistics with multiple comparisons by Bonferroni correction. (**I**) Enrichment (log_2_ fold enrichment) of NDD risk genes among DEGs induced by each perturbation, plotted against -log_10_(BH-adjusted p-value). Point size indicates total DEG count for the target. Labeled points indicate top 10 perturbations by fold enrichment or p-value. (**J**) STRING protein-protein interaction network of pooled DEGs from NDD-enriched perturbations, showing functional convergence on RNA processing and synaptic pathways(minimum interaction confidence score 0.7). Nodes represent proteins and edges represent known or predicted functional associations. Clusters derived from MCL clustering with inflation parameter 1.3. Proteins in bold font are encoded by NDD genes (N = 373)^49^.

To determine whether perturbation of these disease-associated and functionally defined gene sets produced stronger transcriptional burden in specific neuronal populations, we compared the total number of DEGs under each perturbation subset (Fig. 4B; Table S9). ASD- and DD-associated perturbations induced robust transcriptional changes in cortical neurons, spanning both upper- and deep-layer glutamatergic populations. Effects were also observed in thalamic, hypothalamic, and midbrain neurons, whereas changes were relatively weaker in the dentate gyrus of the hippocampus. Psychiatric disorder genes showed comparatively modest signals, potentially reflecting the young age of the animals assayed in this study (P45). Alzheimer’s disease (AD) genes produced minimal changes, consistent with the neuronal focus of this screen and the expected glial contributions to AD pathogenesis^55^. As expected, LoF-constrained genes produced widespread effects across cell types, whereas LoF-tolerant genes showed clean baseline responses. Genes previously classified as essential^57^ also showed clear signals – establishing in vivo essentiality in postmitotic neurons with cell-type resolution.

These patterns were corroborated at the level of individual perturbations. ASD/DD gene perturbations, essential gene perturbations, and LoF-constrained gene perturbations all produced more total DEGs than baseline (Fig. 4C). Perturbation of genes associated with dominance inheritance produced significantly greater DEG burden compared to lower-ranked genes, whereas recessive and X-linked categories showed no significant difference (Fig. 4D). This aligns with our expectation that dosage-sensitive genetic perturbations exert stronger downstream transcriptional effects. Among ASD and NDD gene perturbations, those with stronger de novo association signals, reflected by lower false discovery rate (FDR) values estimated using the Transmission And De novo Association (TADA) model, produced significantly more DEGs (Fig. 4E), indicating that our transcriptomic readout scales with human genetic traits and effect sizes.

Given that NDD gene perturbations with the lowest FDR values produced the greatest transcriptional impact, we further asked whether their effects were uniformly distributed across cell types or exhibited cell-type specificity. Among the 73 significant NDD genes (FDR = 0) with the largest transcriptional effects, DEG distributions across cell types revealed cell-type specificity, with thalamic populations showing particularly large effects (Fig. 4F-G, S11A; Table S9). For example, perturbation of *Stxbp1*, a synaptic vesicle release gene, produced pronounced transcriptional changes in cortical L4/5 IT neurons, including downregulation of synaptic genes such as *Nrxn2* and *Grin2b*. In contrast, perturbation of *Ctnnb1*, a key effector of Wnt signaling, produced stronger effects in thalamic neurons, including dysregulation of transcriptional regulators and signaling components. These examples illustrate how distinct disease genes perturb different neuronal populations and molecular pathways, an architecture that becomes visible only at the scale and cellular resolution of this screen.

We next assessed whether the downstream transcriptional changes linked to each perturbation aligned with disorder-associated molecular signatures across neuronal populations (Fig. 4H; Table S9). Consistent with the DEG burden analysis (Fig. 4B), known ASD- and DD- risk genes were preferentially enriched among perturbation-induced DEGs in neuronal populations that exhibited stronger transcriptional responses to ASD- and NDD-linked perturbations. Psychiatric and neurodegenerative disorders, which showed overall weaker DEG burden, also exhibited more modest enrichment of known risk genes. Together, these results suggest that the observed transcriptional patterns are not random consequences of gene disruption but reflect biologically coherent disease-related molecular signatures.

To extend this analysis from the cell-type level to individual perturbation targets, we further asked whether DEGs induced by any single perturbation were themselves enriched for 373 NDD risk genes (FDR ≤ 0.05)^49^ across cell types (Fig. 4I, S11B-C; Table S9). Although perturbations generating the largest total number of DEGs (e.g., *Tpr*, *Psmb4*) showed enrichment largely as a function of scale, the strongest effect sizes were observed among perturbations producing more selective transcriptional responses. Notably, the highest fold enrichment was observed for *Ankrd11*, itself a known NDD gene. Overall, many perturbations showed enrichment for NDD genes, consistent in part with target selection strategy and the relatively high baseline expression of NDD genes in brain-derived cell types.

As highly expressed genes are more likely to be detected as DEGs, we tested whether NDD enrichment reflected baseline expression bias. We generated 1,000 bootstrapped non-NDD gene sets matched to the NDD genes by whole-brain expression distribution and recalculated enrichment for each target. In many cases, expression-matched control genes exhibited enrichment comparable to or exceeding that of NDD genes, indicating that baseline expression accounts for a substantial component of the observed signal (Fig. S11B). However, a restricted subset of targets retained NDD fold enrichment exceeding the matched background, consistent with selective overrepresentation beyond baseline expression effects (Fig. S11B).

We therefore focused on the five targets with the strongest NDD enrichment exceeding median expression-matched controls (*Slc17a5*, *Ankrd11*, *Mdh1*, *Xrn1*, and *Gatad1*). Pooling their non-self DEGs and constructing a high-confidence STRING protein-protein interaction network revealed significant connectivity and clustering into shared functional modules (Fig. 4J; Table S9). Although modules related to proteostasis and mRNA splicing were present, consistent with broader perturbation effects across the dataset, synaptic signaling emerged as the dominant organizing feature of the NDD-enriched network (Fig. 4J). Multiple synaptic modules were represented, including postsynaptic GPCR signaling, membrane ion homeostasis (Na⁺/K⁺ and SERCA), Kv3 voltage-gated potassium channels, and calcineurin/PP2A-mediated pathways, but convergence across targets was most pronounced within the NMDA receptor and postsynaptic signaling cluster (Fig. S11C). Splicing and ubiquitination likely reflect general transcriptional responses to perturbation rather than NDD-specific biology. The consistent overlap of target DEG lists within the NMDA-centered synaptic module indicates that excitatory postsynaptic signaling represents a shared axis of vulnerability across genetically distinct NDD-associated perturbations (Fig. S11C).

Our results uncover cell-type-specific and pathway-level signatures that provide insight into how human disease genes disrupt brain-wide molecular and transcriptional programs. These results demonstrate that our genome-scale in vivo Perturb-seq approach provides a scalable and interpretable framework for probing the biological mechanisms of risk mutations in their disease-relevant cellular contexts.

## Discussion

This study demonstrates that massively parallel genetic perturbation profiling in the intact brain can resolve the functional architecture of neuronal gene regulation at single-cell resolution. Scaling Perturb-seq to nearly 2,000 genes across the mouse brain was enabled by our optimized in vivo delivery^7,8^, improved gRNA recovery through molecular vector designs^9^, and advances in combinatorial indexing-based single-cell sequencing^58^, building on recent efforts to expand the throughput of perturbation screens and the ‘virtual cell’ efforts^59-65^. Efficient data exploration and processing of the resulting datasets was enabled by GPU-accelerated analysis pipelines leveraging rapids-singlecell, underscoring that continued investment in scalable computational tools will be essential as perturbation atlases grow in scope^66^.

A central finding is that the transcriptional consequences of many disease-relevant genes in the brain are remarkably sparse and cell-type-specific – a level of resolution that is largely invisible to bulk tissue profiling. Disease-associated genes, particularly those linked to NDD/ASD, produced transcriptional effects restricted to defined neuronal populations, and perturbations of distinct risk genes frequently converged on shared downstream programs^67^. This convergent architecture is encouraging from a therapeutic perspective: if multiple genetic etiologies funnel through common cell-type-specific regulatory programs, then targeting those shared nodes could, in principle, address multiple causal mutations – a framework for genome-informed therapeutic design that warrants direct functional testing in the near future^68^. A natural next step is to ask whether perturbing convergent targets can rescue or reverse the phenotypic effects of individual risk-gene mutations, and to extend this approach to additional organs and model systems where the same scalable delivery and sequencing framework can be applied.

Several caveats should be noted. This current screen is limited to loss-of-function perturbations at a single timepoint, using only RNA-seq readouts, and excludes early developmental windows, gain-of-function effects, and non-cell-autonomous interactions. AAV9 tropism further restricts access to certain populations, notably microglia^69^. Cell coverage per perturbation, while sufficient to detect strong transcriptional effects using conventional differential expression approaches such as the Wilcoxon test, limits power for detecting subtle or rare phenotypes. Emerging machine-learning–based approaches for modeling single-cell perturbation responses may improve sensitivity and interpretability of perturbation effects and enable robust phenotype detection with substantially fewer cells in future studies. Despite these constraints, this dataset reveals core biology and cell-type-resolved insights that motivate broader application of in vivo perturbation profiling across the nervous system and beyond.

## Acknowledgments

We thank Taurean Dyer and Gary Burnett for their advice and support on the compute infrastructure; Paul Lund and Peter Smibert for technical advice on the FLEX snRNA-seq library construction optimization; Tom Nowakowski for advice on the manuscript; Scripps DAR and Flow Cytometry Core for assistance; and all members of the Jin lab for their support.

## Data sharing

Data generated for this study are available through the Hugging Face repository (https://huggingface.co/datasets/perturbai/wholebrain_crispr_atlas) and the UCSC Cell Browser (https://cells.ucsc.edu/?ds=whole-brain-perturb). The analysis pipeline is deposited on GitHub repository (https://github.com/jinlabneurogenomics/wholebrainperturbseq).

## Materials and Methods

### Animals

All animal experiments were performed according to protocols approved by the Institutional Animal Care and Use Committees (IACUC) of The Scripps Research Institute. Retro-orbital injection of pooled AAV library was performed in P16 C57BL/6J or CD-1 x Cas9 transgenic mice^7^ of varying sexes and weight across litters. Brains were harvested 3-4 weeks post-inject between P37 and P44 for single-nucleus and immunohistochemistry experiments (Table S3). All mice were kept in standard conditions (a 12-h light/dark cycle with ad libitum access to food and water).

### Cell culture

Mammalian cell culture experiments were performed in the HT-22 mouse hippocampal neuronal cell line (Millipore Sigma, #SCC129) grown in DMEM (Thermo Fisher Scientific, #11965092) with 25mM high glucose, 1mM sodium pyruvate and 4mM L-Glutamine (Thermo Fisher Scientific, #11995073), additionally supplemented with 1x penicillin–streptomycin (Thermo Fisher Scientific, #15140122), and 5-10% fetal bovine serum (Thermo Fisher Scientific, #16000069). HT-22 cells were maintained at confluency below 80%.

### AAV vector construction and production

Viral vector used in this study was previously reported^9^ (Addgene #TBD). Guide RNAs used in this study were designed using CRISPick^70,71^ and top 4 guides for each target gene were selected. Full sequences of gRNAs used in this study are listed in Table S1. Pooled gRNA oligonucleotides were cloned in this vector as described in Joung et al. 2017^72^. Briefly, oligonucleotides encoding gRNA sequences with flanking sequences homologous to the vector backbone were produced (Twist Biosciences) and amplified using primers YZJ_019 and YZJ_024 (Table S10). They were then inserted into LguI (Thermo Scientific, ER1931) digested vector backbone using Gibson Assembly (NEB E2611). Assembled DNA was then electroporated in Endura electrocompetent cells (Lucigen, #60242-1) using BTX ECM 630 electroporation system (Harvard Apparatus, #45-0051). Presence and distribution of each gRNA were verified by PCR pooled plasmid library to attach Illumina sequencing primers (YZJ_025-043, Table S10), followed by sequencing using Illumina iSeq 100.

AAV production and titration were adapted from Challis et al. 2019^73^. Briefly, HEK293T cells were transfected with above pooled plasmid library or plasmid encoding a PiggyBac transposase^7^, along with AAV-PHP.eB capsid (Addgene #103005) and pHelper plasmids using PEI (Polysciences, #24765-1) at 80-90% confluency. After 1 media change at 20 hours post-transfection, viral supernatant was collected at 72 hours and 120 hours post-transfection. Transfected cells were also harvested and digested at 120 hours post-transfection. Viral particles in the collected media were pelleted using PEG at the final concentration of 8% wt/vol (Fisher Scientific, #BP233100) and purified via ultracentrifuging (Beckman Coulter, #A94516) at 59,000 rpm for 2 hrs and 25 min at 18 °C. Purified AAVs were then concentrated in DPBS with 0.001% Pluronic F-68 using Amicon® Ultra Centrifugal Filter 100kDa (Millipore Sigma, #UFC910024). AAV particles were then digested and titrated using primers (Table S10) targeting WPRE within the packaged transgene. Presence and distribution of each gRNA were then again verified by PCR on a pooled AAV library to attach Illumina sequencing primers (YZJ_025-043, Table S10), followed by sequencing on an Illumina iSeq 100.

### Selection of 1,947 gene targets

To establish a brain-wide functional genomics atlas, we curated a target gene library prioritized by human disease relevance, transcriptional regulatory function, and central nervous system (CNS) expression. **Gene Selection Criteria and Scoring Rubric:** Candidate genes were restricted to protein-coding loci situated on chromosomes 1-21, X, and Y, as defined by the MGI GRCm39 reference. To ensure biological relevance to the brain, we excluded ribosomal protein genes and focused on those with established expression within neural tissue. A multi-tiered scoring system was implemented to prioritize genes based on their association with four primary Disease Ontology (DO) categories: Genetic Disease (DOID:630), Diseases of Mental Health (DOID:150), Diseases of Metabolism (DOID:0014667), and Central Nervous System Disease (DOID:331). Genes were further filtered by functional class, specifically targeting Transcription Factors (GO:0000981) and Signaling Receptors (GO:0038023). We also integrated a manually curated set of high-confidence NDD risk genes. **Quantitative Ranking and Expression Weighting:** To rank candidates, we developed a weighted scoring algorithm. Baseline expression data were derived from a comprehensive mouse brain cell atlas^74^, where the mean expression for each neuronal gene was calculated and ranked into deciles (percentiles 1–10). The final selection score was calculated as follows: S = 10 x w_c + 2 x (w_a + w_b) + 0.2 x Ep, where: w_c represents primary disease association weight, w_a and w_b represent secondary functional and ontological weights, and E_p represents the expression percentile (decile) ranking. Applying a score cutoff of 3.8, we identified a final library of 1,948 target genes. **sgRNA design:** We designed four gRNAs per gene by targeting coding exons near the 5’ end of the transcripts along with 400 non-targeting control guides and 400 safe-targeting (ST) gRNAs targeting 100 intronic and intergenic regions.

### Genomic (insertion-deletion) analysis to evaluate in vitro gRNA activities

HT-22 mouse hippocampal neuronal cells (Millipore Sigma, #SCC129) were plated in 96-well plates at 15,000 cells per well 24 hours prior to transfection to achieve 60-80% confluency the next day. 200 ng of plasmids encoding each individual single gRNA expression were separately transfected into each well using lipofectamine 3000 (Thermo Fisher #L3000001). Transfection media were replaced with fresh growth media 24 hours post-transfection. Genomic DNA was extracted from each well 48 hours post-transfection using QuickExtract DNA extraction solution (Lucigen #QE09050). The genomic regions of each targeted locus were PCR amplified and attached with Illumina sequencing adaptors (Table S10) for downstream sequencing with Illumina iSeq 100. Fastq files from sequencing were aligned to amplicons and insertion/deletion analyzed using CRISPResso2 package^75^.

### Viral vector administration

All mice in this study were injected retro-orbitally with AAVs (30-50 μL per animal). C57BL/6J mice were injected at P16 at 1e^9^, 6e^8^, or 1.5e^8^ units per gram body weight of pooled AAV library, along with 1e^11^ units of PiggyBac transposase AAV, and brains were harvested between P37 and P44 for immunohistochemistry. Cas9 transgenic mice were injected at P16 at 3-6e^8^ units per gram body weight of pooled AAV library, along with 1e^11^ units of PiggyBac transposase AAV, and brains were harvested between P37 and P44 for single-cell experiments.

### Immunohistochemistry

Mice were anesthetized and transcardially perfused with ice-cold PBS followed by ice-cold 4% paraformaldehyde in PBS. Dissected brains were postfixed overnight in 4% paraformaldehyde at 4 °C, and then washed in PBS. The brains were embedded in 2% agarose and sectioned at 50 μm using a vibratome. The slides with tissue sections were incubated with blocking media (10% donkey serum (Sigma Aldrich, #S30–100ML) in 0.3% Triton with PBS) for 2 hrs, then incubated with primary antibodies diluted in blocking media overnight at 4 °C. Slides were washed with PBS with 0.3% Triton 6 times to remove the excess primary antibody. Secondary antibodies were applied at 1:1000 dilution in blocking media and incubated for 2hr at room temperature. Slides were then washed 4 times with PBS with 0.3% Triton, incubated with DAPI for 10 min, and washed twice with PBS, before mounting with Antifade Mounting Medium (Vector Laboratories, #H-1700–10). The primary antibodies and dilutions were: Chicken anti-GFP antibody (ab16901, 1:500; Millipore), Rabbit anti-RFP antibody (600-401-379, 1:1000; Rockland). All images were taken using a Nikon AX Confocal Microscope with a 10x air or 20x air objective. Cell numbers were quantified using CellProfiler.

### Nuclei isolation and FACS-enrichment

Mice between the ages of P37 and P44 were anesthetized with isoflurane, disinfected with 70% ethanol and decapitated. Brains were quickly extracted and divided into two hemispheres. Each hemisphere was diced into ∼3-5 mm cubes, placed in Eppendorf tubes, and flash frozen in liquid nitrogen. Nuclei isolation buffer was prepared with 250 mM sucrose (Sigma #84097), 25 mM KCl (ThermoFisher #AM9640G), 10 mM HEPES (ThermoFisher #15630080), and 5 mM MgCl2 (Sigma #M1028). Each individual hemisphere was placed in dounce grinders (Sigma D9938) filled with 10 mL cold lysis buffer (nuclei isolation buffer containing 1 mM dithiothreitol (Sigma #646563), 0.1% NP-40 substitute (Fisher #50-220-4931), 2% BSA (Miltenyi #130-091-376), 10mg/mL Kollidon VA64 (Sigma #190845), and 0.2 U/μL RNase Inhibitor (Fisher #NC1081844)). 7 mL more of cold homogenization buffer was added to the douncer and the tissue was allowed to lyse on ice for 3 min. The homogenized and lysed tissue was then transferred through a 70 μm strainer (VWR #21008-952) to a tube pre-filled with 10 mL of cold wash buffer (nuclei isolation buffer with 1 mM dithiothreitol (Sigma #646563), 2% BSA (Miltenyi #130-091-376), and 0.2 U/μL RNase Inhibitor (Fisher #NC1081844)). The sample was then centrifuged at 300xg for 10 min, and the pellet was resuspended in 6 mL wash buffer, filtered through a 40 μm strainer (Fisher #07001106), and stained with 1:1000 dilution of Alexa Fluor 647 Anti-NeuN antibody (Abcam #ab190565) for 15 min on ice. The stained sample was then washed twice with cold wash buffer, followed by centrifuging at 250xg and 200xg at 4 °C for 8 min each. The pellet was then resuspended in 5mL 1x Fix & Perm Buffer (10x Genomics, PN-2000517) with 4% paraformaldehyde and fixed overnight at 4 °C. Fixed sample was then quenched with Additive C (10x Genomics, PN-2001332) and Quenching Buffer (10x Genomics, PN-2001300). GFP^+^ and NeuN^+^ nuclei were collected into cold PBS with 1% BSA and 0.2 U/μL RNase Inhibitor (Fisher #NC1081844) using the MA900 (Sony), MoFlo Astrios (Beckman), or Bigfoot Spectral Cell sorter (Thermo Fisher) with 100 μm sorting chips. Sorted nuclei were then resuspended in 1x Quenching buffer (10x Genomics, PN-2001300) with 10% Enhancer (10x Genomics, PN-2000482) and 10% glycerol (Millipore Sigma #G5516-100ML) and banked at -80 °C for storage.

### Single-nucleus RNA-seq library preparation and processing

Single-nucleus RNA-seq libraries were constructed using the Chromium GEM-X Flex Apex/v2 platform (10x Genomics). Each FACS-sorted hemisphere sample was hybridized separately using mouse whole transcriptome probes (10x Genomics, PN-1000931) and custom-designed probes that targeted each of the CRISPR gRNAs (Table S1), using a temperature ramp from 60 °C to 42 °C and held at 42 °C overnight. Each hybridized sample was then subpooled with different sample barcodes at 25,000 nuclei per barcode and incubated at 42 °C for 2 hrs. Barcoded samples were pooled, washed, and loaded into each 10x GEM channel targeting recovery at maximum 1 million nuclei per channel. The gene expression and CRISPR libraries were then prepared following manufacturer’s protocol and sequenced with Illumina Novaseq X Plus 25B 100-cycle flowcells (Read 1: 54 bases; Read 2: 50 bases), with sequencing coverage greater than 20,000 reads per cell for gene expression libraries and 3,000 reads per cell for CRISPR libraries. BCL files were converted to FASTQ files using BCLconvert with default parameters. The “cellranger multi” command (cellranger v10.0.0) was used to align FASTQ reads to Mouse Transcriptome Probe Set v2.0 (10x Genomics) and a custom feature reference containing the gRNA sequences used in this study (Table S1).

### Cell-level QC and cell type annotation

Several processing analyses were applied to filter cell barcodes before downstream analyses. To remove potentially dying cells, cell barcodes with fewer than 2000 detected genes were removed. Each 10x channel was processed with the scDblFinder R package (v1.24.0) with the parameter dbr=0.15. To limit excessive memory use, for channels with more than 200k cell barcodes, cells were randomly split into 100k ‘samples’ and scDblFinder was run using multiSampleMode = ‘split’. Only barcodes designated as ‘singlet’ were retained for downstream analysis.

Additionally, we identified a group of cells with an ambiguous cell type and statistically indistinguishable expression from the average expression across all cells. To identify these, we created an “ambient-score” statistic that measured, for each cell, its mean squared difference from the ambient background, and we removed cells with a threshold on this score. In detail, we define:

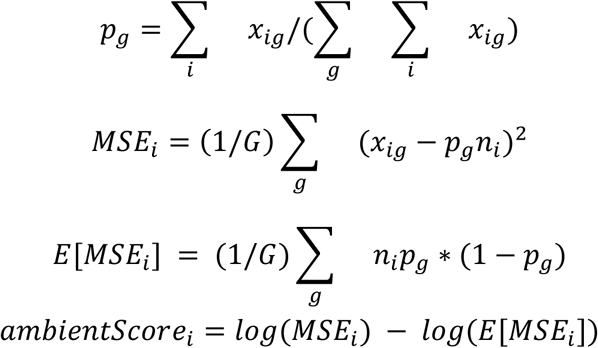

Where x_ig_ is the UMI count for gene g in cell i, G is the total number of genes in the flex reference, and n_i_ is the number of UMIs detected for cell i. p_g_ represents the proportion of detects for gene g, MSE_i_ is the mean squared difference of the detection of each gene in cell i from a random draw of n_i_ UMIs from the background, and E[MSE_i_] is the expected MSE given a depth of n_i_. The empty droplet cells formed a distinct peak at ambientScore=0, and were removed with a threshold of 0.09 (4.3% of cells).

A final clustering step was used to filter cells for downstream analyses. Cells were embedded into 100 dimensions using scVI (scvi-tools v1.4.1) and clustered using the Leiden algorithm with resolution 1.0 as implemented in scanpy (v1.12). A small subset of clusters (clusters 1, 2, 3, 6, and 17, consisting of 2.3% of cells) showed evidence of being residual droplet multiplets not detected by scDblFinder: ambiguous neuron class from MapMyCells, and a mixture of markers from diverse neuron and non-neuron cell types. These were removed from downstream analysis along with clusters 57 and 83 which corresponded to astrocytes and oligodendrocytes.

### MapMyCells fidelity test

Whole brain cell identities were assigned using the Allen Brain Cell (ABC) Atlas’ *cell_type_mapper* package to apply the MapMyCells approach with 10x Whole Mouse Brain RNA-seq (CCN20230722) as reference and hierarchical mapping algorithm. To ensure MapMyCells could faithfully retrieve cell types from mixed brain regions, we performed a validation experiment (Fig. S1): we dissected five major brain regions from two mice (cortex, striatum, hippocampus, midbrain/thalamus/hypothalamus combined, and cerebellum) and constructed snRNA-seq libraries using the Chromium GEM-X Flex v1 platform (10x Genomics) with each brain region from each mouse individually barcoded (Fig. S1E). Each dissected region mostly formed its own clusters, with some formed clusters in their anatomically adjacent regions, likely due to tissue contamination during dissection (Fig. S1F). Additionally, we inferred the developmental origin neighborhood^6^ for each cluster (Fig. S1G) and found nuclei isolated from different brain regions were largely represented by their corresponding developmental neighborhoods with little to no contamination from developmentally and anatomically distant brain regions (Fig. S1H). This indicates that MapMyCells can assign brain cell types with high fidelity and can be applied to a dataset containing cell types from across the entire brain.

### gRNA filtering and perturbation assignment

To improve the accuracy of guide assignment, we used two filtering stages to remove guide UMIs that likely arose due to sequencing or PCR errors. First, we applied a dynamic read-support threshold for guide UMIs. The threshold was defined as the maximum integer k such that discarding UMIs with at least k reads resulted in a cumulative loss of no more than 1% of total sequenced guide reads. Second, for the remaining UMIs, we merged UMIs that had the same cell and guide barcode, and UMI barcodes within an edit distance of 2. After this filtering process, guides were then assigned to cells if their UMI count in a cell was at least 3.

### UMAP visualization

For global UMAP representation (Fig. 1B), using the filtered post-QC cells annotated using MapMyCells, gene expression counts were normalized using all detected genes and then restricted to 6,966 marker genes derived from a 8,460 marker gene panel^6^. To de-noise expression values and visually cluster groups, we applied iterative k-nearest neighbor (KNN) imputation (k=15, metric=cosine). For each level in the neighborhood/class/subclass hierarchy, we restricted neighbor search for a query cell to reference cells in the same group, and marker gene expression for the query cell was replaced by the mean expression of its neighbors. For the top-level neighborhood, imputation is done with the log-norm X feature space, and all subsequent levels use the imputed X matrix from the preceding round. PCA was then computed on the final imputed matrix, selecting the top 100 components as inputs to the final 2D UMAP embedding (n_neighbors = 25, metric = euclidean, md = 0.4).

For neighborhood UMAP representations (Fig. 1F, S4A), cells were separated by neighborhood and embedded independently. The 100-dimensional scVI latent representation generated during QC was used as input for 2D UMAPs (n_neighbors = 25, metric = euclidean, md = 0.4).

### E-distance

To quantify the similarity of phenotypic shifts induced by different genetic perturbations, we calculated pairwise E-distance (Energy Distance)^76^ between each pair of perturbations, including the control cell population. First, we subset the data to only those conditions that have at least 5 differentially expressed genes with an absolute log fold change greater than 0.5 (|*LFC*| > 0.5). For each cell type sub-population, raw counts were initially normalized to a target sum of 10^4^ (CP10K) and subsequently log-transformed using the formula *log*(1 + *x*). For feature selection, we identified the top 2,000 highly variable genes (HVGs) specific to the cell type. Following this, the data were scaled to unit variance and zero mean, and Principal Component Analysis (PCA) was applied to reduce dimensionality to the top 50 principal components, which capture the primary axes of variance within that cell type.

The E-distance was computed in the 50-dimensional PCA space using the pertpy library (v1.0.5)^77^. For two distributions of cells *X* (gene target *X*) and *Y* (gene target *Y*), the E-distance is defined based on the expected Euclidean distances between cells within and across the two groups. Specifically, it measures whether the mean distance between cells from *X* and *Y* is significantly greater than the mean distances between cells within *X* and within *Y*. The resulting pairwise distances were recorded for every pair of perturbations (*X*, *Y*) within each cell type, providing a standardized metric to quantify the similarity of phenotypic impact across different genetic targets. We repeated these steps for Neighborhood and Whole-Brain cellular resolutions.

### Differential abundance and compositional analysis

#### Library-Normalized Relative Fitness Test (Fisher’s Exact Test)

To determine if specific genetic perturbations induced cell-type-specific toxicity or fitness defects, we performed a Library-Normalized Relative Fitness Test. For each perturbation sgRNA *P* and cell type *C*, we constructed a 2 × 2 contingency table. We compared the observed cell counts with their respective frequencies in the AAV NGS library for perturbed and the Non-targeting Control (NTC) cells.

The contingency table for each test was defined as follows:

- A: Expected count of cells with perturbation sgRNA *P* in cell type *C*, calculated as the total cells in cell type *C* multiplied by the NGS library fraction of *P*.
- B: Observed count of cells with perturbation sgRNA *P* in cell type *C*.
- C: Expected count of NTC cells in cell type *C*, calculated as the total cells in cell type *C* multiplied by the NGS library fraction of the NTC guides.
- D: Observed count of NTC cells in cell type *C*.

Statistical significance was determined using a two-sided Fisher’s Exact Test via scipy library (1.16.3)^78^. Relative fitness was quantified using the odds ratio. Odds ratio > 1.2 indicates a fitness defect (toxicity) where the perturbation is recovered at a lower rate than the NTC within that specific cell type. Multiple testing correction was performed using the Benjamini-Hochberg procedure ^15^ across all perturbation-cell type pairs, with significance defined as an FDR < 0.05.

The analysis was conducted at Neighborhood cell type-resolution. Analysis was performed at the individual guide level. To ensure biological robustness and minimize the impact of stochastic sampling or off-target effects, a genetic perturbation was only considered to have a fitness defect if at least two out of four independent guides met the significance and effect-size criteria.

#### Guide Abundance Modeling (MAGeCK)

As another way to quantify the fitness impact of each genetic perturbation, we employed a "Day 0 vs. Day X" dropout model. The initial budget (Day 0) for each sgRNA was defined by its proportional representation in the NGS library, scaled to the total number of cells captured within each specific cell-type subclass. The observed abundance (Day X) was defined as the raw cell count for each sgRNA within the cell population. Differential abundance analysis was performed using the MAGeCK (v0.5.9)^14^ pipeline. To account for the overdispersion typical of single-cell sequencing data, MAGeCK utilized a Negative Binomial (NB) distribution to model the relationship between the mean and variance of guide counts. For each cell type at the Neighborhood resolution, we compared the observed cell counts against the calculated expected budget. To ensure the robustness of our findings, we utilized the Robust Rank Aggregation (RRA) algorithm. RRA identifies significant gene-level hits by evaluating the rank consistency of all four independent sgRNAs targeting a single gene. The resulting p-values were adjusted for multiple hypothesis testing using the Benjamini-Hochberg (BH) procedure^15^. Cell populations from genetic perturbations were classified as significantly based on a FDR threshold of < 0.05 and a consistent negative LFC of < - 0.25.

### Differential expression (DE) analysis

Differential expression (DE) analysis was performed to identify changes in gene expression associated with specific genetic perturbations across distinct cell types. To ensure the highest fidelity in assigning genotype to phenotype, we restricted our analysis exclusively to cells with a single, unambiguously assigned sgRNA. This inclusion criterion was strictly applied across all cells, including cells containing targeting guides, non-targeting (NT) control guides, and safe-targeting (ST) control guides. To establish a statistically robust baseline, we aggregated cells from all 400 distinct non-targeting guides into a unified control pool. All experimental perturbations, including targeting guides and safe-targeting controls, were statistically compared against this pooled non-targeting population. To empirically determine the FDR, we utilized a library of 400 safe-targeting guides. These 400 safe-targeting guides were randomly partitioned into 100 pseudo-genes, with each pseudo-gene comprising a pool of 4 independent guides. These pseudo-genes were treated as active perturbations and the same DE analysis was performed against the non-targeting pool.

To conduct the differential expression testing, we employed the Wilcoxon rank-sum test approach, implemented using the Scanpy framework^79^ (v1.12). For each cell type, the expression matrix was filtered to include only genes expressed in at least 10 cells within that specific subpopulation. To ensure sufficient statistical power, we considered only perturbations and control groups with at least 20 cells per cell type. For each valid cell-type/perturbation pair, we performed a two-group comparison against the non-targeting control population. The resulting p-values were adjusted for multiple hypothesis testing using the Benjamini-Hochberg (BH) procedure^15^.

### Computing perturbation effect vectors

To compare perturbation effects across all cell types, we took the subset of gene targets with at least 5 differentially expressed genes (Wilcoxon test, FDR < 0.05) in at least one cell type and fit a variational autoencoder^80^ to cells with these perturbations (as well as non-targeting control cells). To ensure that vectors in the models latent space are comparable, we used LDVAE algorithm^81^ which features a linear output decoder, and to focus the model on perturbation effects (and not baseline tissue differences) we added the cell’s cluster (in the original scVI embedding) as a categorical covariate. To isolate perturbation+tissue combinations where an effect was detected, we used an E-Distance test in this latent space, which resulted in 1,218 significant effects (FDR < 0.05).

Summary perturbation-effect vectors were then computed by averaging the latent space embedding of cells sharing the same perturbation target within the same tissue and subtracting out the average of control cells (cells with a single non-targeting control guide) in the same tissue. These vectors were then compared across the full dataset using cosine similarity as a metric. Heatmaps in Fig. 3B, 3D, and S9 were generated applying hierarchical clustering (using euclidean distance) to the cosine similarity matrix.

### Curation of disorder risk genes and functional stratification

Risk genes associated with neurological disorders and conditions were curated from large-scale exome sequencing studies, genome-wide association studies (GWAS), and clinically curated gene panels. For neurodevelopmental diseases, ASD and NDD risk genes were obtained from large-scale exome studies of de novo variants using an FDR threshold of ≤ 0.05^49^. Genes associated with developmental delay were obtained from exome-wide significant genes reported in the Deciphering Developmental Disorders (DDD) study^50^. Intellectual disability, epilepsy, and ataxia gene sets were curated from the Genomics England PanelApp database, including genes classified as “Green” (high confidence)^51^. Schizophrenia risk genes were obtained from large-scale rare variant exome studies reporting exome-wide significant associations^40^. For psychiatric and neurodegenerative disorders, risk genes were curated from large-scale GWAS meta-analyses, including bipolar disorder (BPD)^52^, major depressive disorder (MDD)^53^, attention-deficit/hyperactivity disorder (ADHD)^54^, and Alzheimer’s disease (AD)^55^. Parkinson’s disease (PD) and Amyotrophic lateral sclerosis (ALS) were curated from the Genomics England PanelApp database. PD genes were obtained from the Parkinson Disease and Complex Parkinsonism panel (version 1.1.28), and ALS genes from Amyotrophic Lateral Sclerosis/motor neuron disease (version 1.74).

Genes were further stratified based on the loss-of-function observed/expected upper bound fraction (LOEUF) score, which quantifies gene tolerance to functional variation^56^. Genes were partitioned into deciles (0 to 9) based on their relative LOEUF rank across all genes. We defined LoF-tolerant genes as those in deciles 6-9, LoF-intermediate genes as those in deciles 3-5, and LoF-constrained genes as those in deciles 0-2. Gene essentiality was defined based on genome-scale CRISPR knockout screens in human cell lines (HAP1 and KBM7)^57^. Curated gene sets were converted to mouse homolog genes using the BioMart homology database^82^.

### Cell-type-specific DEG burden across disorder-associated gene perturbations

The number of DEGs (FDR < 0.05) was quantified for each perturbation across cell types. Perturbations were grouped according to curated genetic categories, including neurodevelopmental, psychiatric, and neurodegenerative disease risk genes, as well as loss-of-function (LoF) constraint groups and essential genes. For each cell type and genetic category, total DEG counts were summed across perturbations targeting genes within that category. DEG counts were log₁₀-transformed for visualization. Heatmaps were generated with cell types ordered by anatomical region and categories grouped by functional class.

### Statistical comparison of DEG counts across groups

The total number of DEGs was statistically compared across defined perturbation groups, including ASD- and developmental delay-associated gene perturbations. For binary comparisons between essential and non-essential genes, differences were assessed using two-sided Wilcoxon rank-sum tests. For genes stratified by LoF tolerance, overall differences were evaluated using the Kruskal-Wallis test, followed by pairwise Wilcoxon rank-sum tests comparing LoF-tolerant and LoF-constrained groups to the LoF-intermediate group.

To examine DEG burden in relation to genetic inheritance mechanisms, inheritance-specific scores (dominant, recessive, and X-linked) were computed by aggregating evidence from exome sequencing datasets and curated knowledge bases. Genes meeting exome-wide significance were assigned 10 points, and those meeting an FDR threshold were assigned 5 points. For LOEUF, genes in the most constrained deciles were assigned higher weights (e.g., 10 points for decile 0), with intermediate deciles assigned proportionally lower scores. Evidence from curated databases (e.g., SFARI, PanelApp, GenCC, Gene2Phenotype, OMIM) was converted into weighted scores based on the reported strength of association (e.g., definitive/strong evidence receiving higher points than moderate/limited evidence). Scores were allocated to dominant, recessive, or X-linked categories according to reported or inferred mode of inheritance and summed to generate consensus scores.

Genes with a dominance score > 20 and a recessive score > 10 were included in the comparison and stratified into top and bottom groups using a median split of the respective score distributions. Comparison by X-linked score was performed using genes located on the X chromosome. Differences between the top and bottom groups were evaluated using the Wilcoxon rank-sum test. For comparisons across perturbations of ASD and NDD risk genes by the level of significance, genes were stratified into five quantiles based on the -log_10_(FDR) distribution.

### Functional characterization of cell type-specific effects of NDD perturbations

NDD genes with TADA FDR = 0 were selected based on prior genetic evidence^49^ and showing high transcriptional effect. For the perturbation of these genes, DEGs (FDR < 0.1) were aggregated across perturbations and cell types to construct a cell type × gene matrix showing DEG counts. For each perturbation, counts were summed within cell types, and perturbations with fewer than 10 total DEGs across all cell types were excluded to focus on robust effects. The resulting matrix was normalized per gene (0-1 scaling) for comparison of relative enrichment patterns across cell types. Total cell numbers per cell type were visualized in a heatmap generated using the pheatmap package. For functional interpretation of *Stxbp1* and *Ctnnb1* perturbation effects, Gene Ontology enrichment analysis was performed using Enrichr^83^. Redundant GO terms were reduced using semantic similarity clustering implemented in rrvgo package (Rel method, similarity threshold = 0.7) ^84^, with representative terms selected based on enrichment significance.

### Cell type-specific enrichment analysis of DEGs with disorder-associated gene sets

The full list of disorder-associated genes, loss-of-function (LoF) constraint categories, and essential genes was subjected to enrichment tests with DEG sets aggregated per cell type with a threshold of FDR < 0.05 and absolute LFC > 0.5 using one-sided Fisher’s exact test (alternative = “greater”). The background was defined as all genes detected in the dataset. P-values were adjusted using Bonferroni correction across all comparisons.

Odds ratios were log2-transformed and capped to limit extreme values for visualization. Enrichment results were displayed as heatmaps, with significant associations indicated by overlaid markers.

### Enrichment of NDD genes across perturbations and network convergence

To assess enrichment of NDD genes among perturbation-induced transcriptional changes, downstream DEGs (FDR < 0.05) were tested for overrepresentation of 373 NDD risk genes (FDR ≤ 0.001) using Fisher’s exact test, with Benjamini-Hochberg correction across targets. Only perturbations with ≥20 unique DEGs were analyzed, and the gene universe comprised all genes tested for differential expression (N = 17,994). Fold enrichment (FE) was calculated as the observed overlap divided by the expected overlap based on the background frequency of NDD genes in the universe.

To control for expression-related bias, we generated 1,000 bootstrapped non-NDD gene sets matched to the NDD genes by whole-brain expression quantiles and computed per-target FE distributions to derive median values and 95% empirical confidence intervals. The top five targets were selected based on BH-adjusted significance (q < 0.05), effect size (log_2_(FE)), and NDD enrichment exceeding the median expression-matched background. For these targets, pooled non-self DEGs were analyzed in STRING (minimum interaction confidence score 0.7) to construct a protein–protein interaction network^85^. Network clusters were identified using Markov clustering (MCL; inflation = 1.3), and connectivity significance was evaluated using the STRING PPI enrichment test. Module convergence was quantified as the percent of genes within each STRING cluster represented in each target’s DEG list.

---

**Figure S1.**
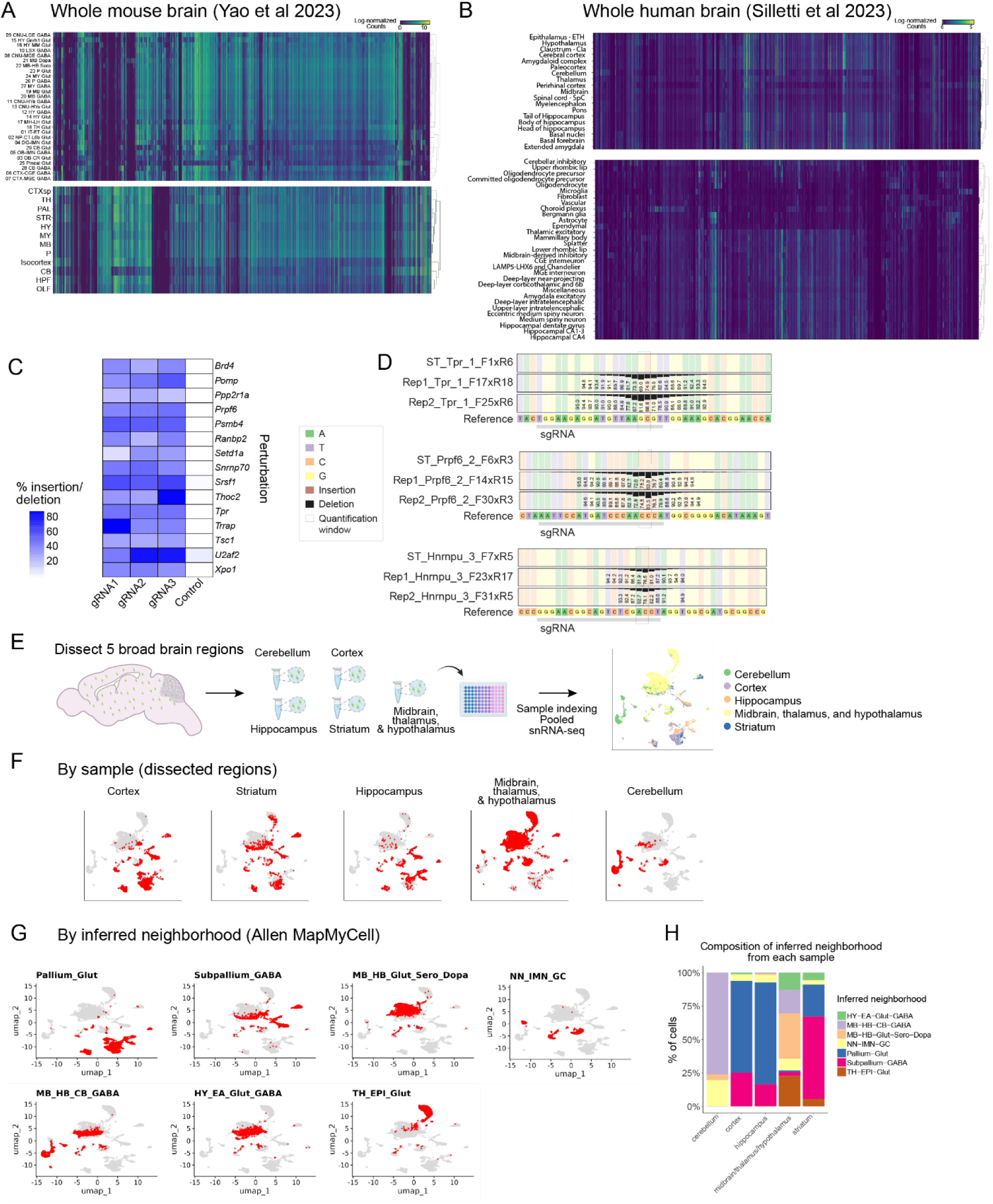
Expression pattern of 1,947 risk genes, in vitro gRNA activity test, and MapMyCells fidelity test. **(A)** Heatmap showing the gene expression levels of 1,947 neurodevelopmental disease-associated risk genes perturbed in this study across different cell types using whole mouse brain atlas^6^. **(B)** Heatmap showing the gene expression levels of 1,947 neurodevelopmental disease-associated risk genes perturbed in this study across different cell types using adult human brain atlas^2^. **(C)** Heatmap of in vitro gRNA activity of 45 selected gRNAs (15 genes, 3 gRNAs per gene) compared to safe-targeting controls by insertion-deletion analysis. **(D)** Examples of nucleotide composition and indel frequency near gRNA cut sites in selected guides (Tpr_1, Prpf6_2, Hnrnpu_3). **(E)** Schematic of snRNA-seq experiment profiling separate brain regions to test MapMyCell^86^ cell type assignment fidelity. **(F-G)** UMAPs of a single-nucleus Flex RNA-seq library collected from whole mouse brains with each major brain region dissected and nuclei extracted separately. Each dissection region (F) and inferred neighborhood (G) are highlighted. **(H)** Stacked bar plot showing percentage of cells from inferred neighborhood recovered from each dissection region.

**Figure S2.**
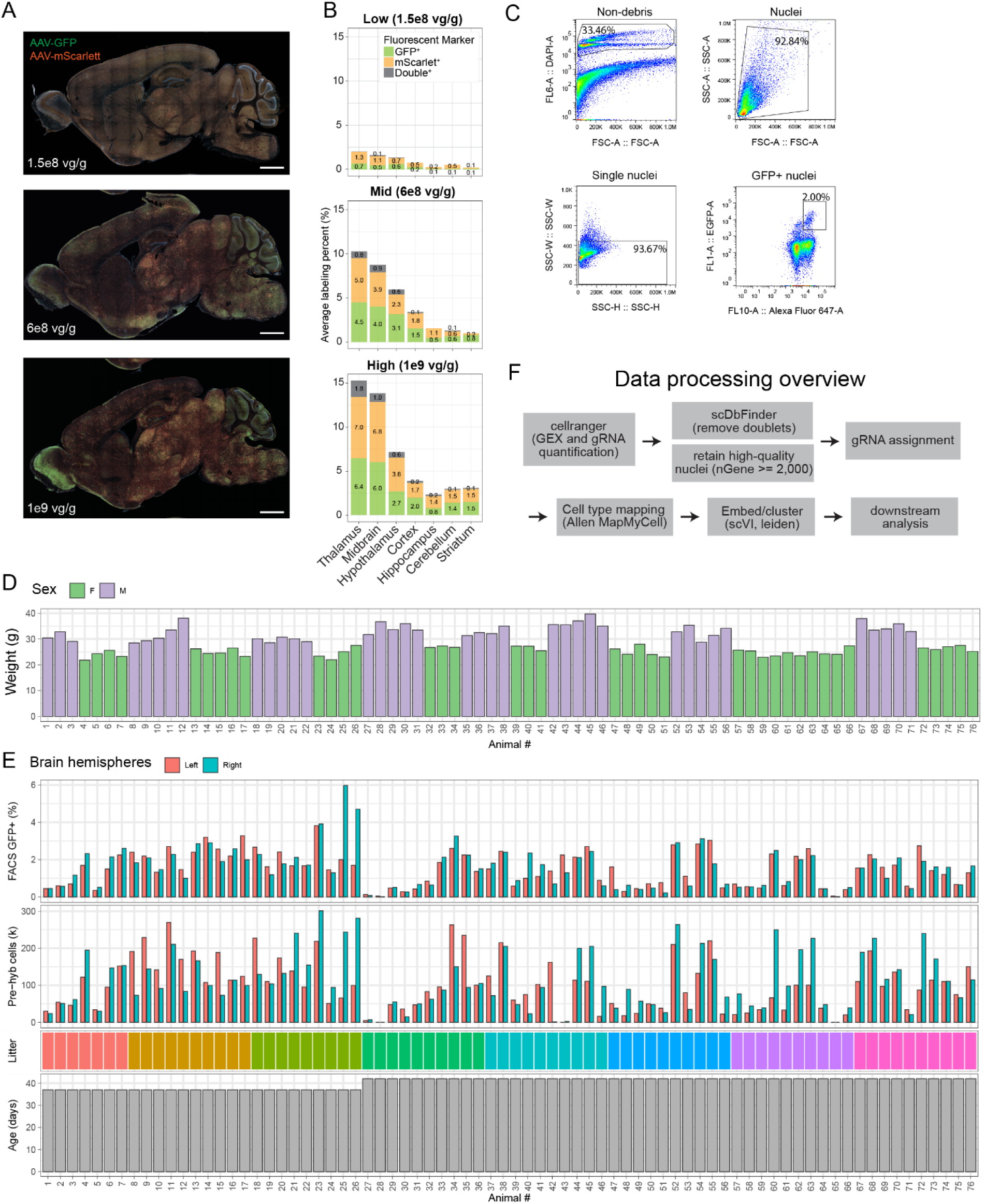
in vivo AAV titer optimization, snRNA-seq data processing, and animal-level metadata. **(A)** Immunofluorescence image of sagittal section of a P37 mouse brain retro-orbitally injected at P16 with high (1e9), mid (6e8), or low (1.5e8) total vg per gram of body weight of AAV PHP.eB encoding either GFP or mScarlet (1:1 ratio) (scale bar = 1 mm). **(B)** Quantification of GFP and mScarlet viral labeling efficiency as well as double labeling rate in (A). **(C)** Representative FACS gating strategy to enrich transduced neuronal nuclei. **(D)** Bar plot of sex and weight at harvest of animals used in this study. **(E)** Animal tracking information showing the litter, age at harvest for each animal, as well as AAV-labeling rate by FACS and total nuclei number per hemisphere used for Flex hybridization. **(F)** Schematic of snRNA-seq data processing and quality control workflow.

**Figure S3.**
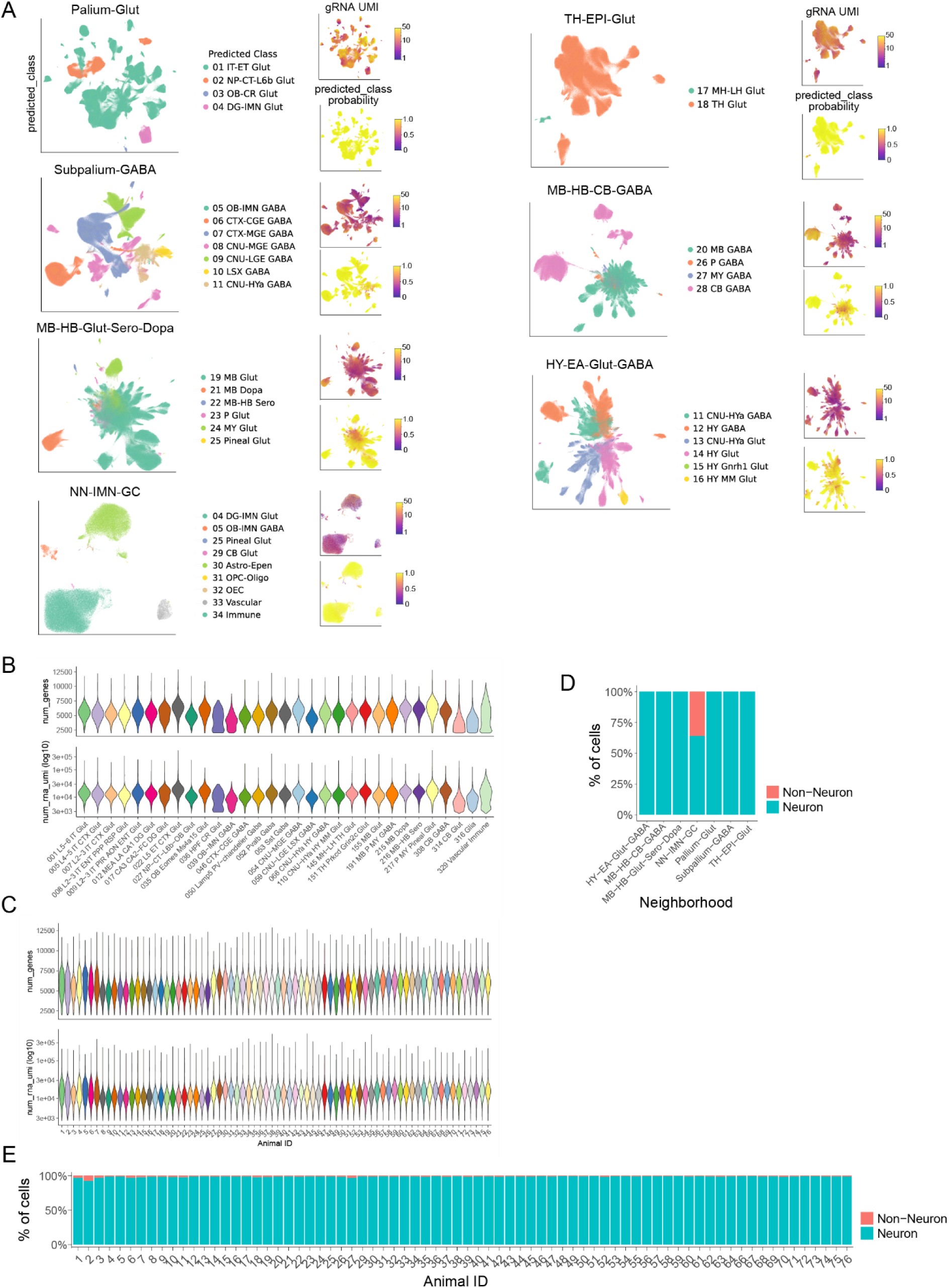
Additional visualization and quality control of snRNA-seq data. **(A)** UMAPs of whole brain in vivo Perturb-seq dataset separated by neighborhoods, colored by inferred cell class and predicted class probability using MapMyCells^86^, as well as number gRNA UMIs per nucleus. **(B)** Violin plots of the number of genes and RNA UMIs recovered per nucleus from each binned cell type. **(C)** Violin plots of the number of genes and RNA UMIs recovered per nucleus from each animal. **(D)** Bar plot of % neuronal versus non-neuronal population recovered from each developmental neighborhood. **(E)** Bar plots of % neuronal versus non-neuronal population recovered from each animal.

**Figure S4.**
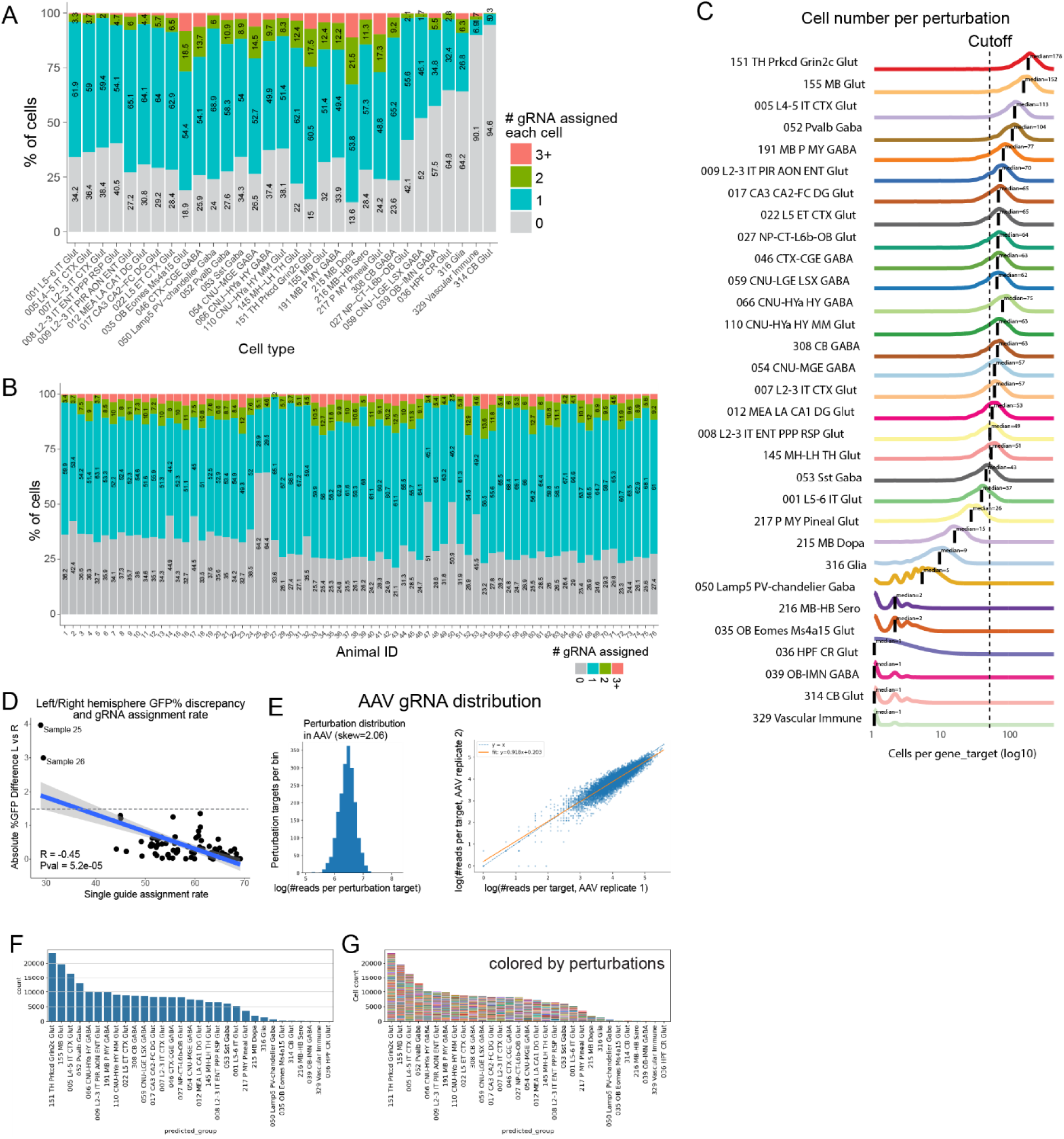
gRNA and gRNA assignment quality control. **(A-B)** Stacked bar plots of percentage of nuclei with no guide, single, double, or multiple guide assignment from each binned cell type (A) and animal (B). **(C)** Density curves of nuclei number recovered per perturbation separated by cell types. Vertical dashed line indicates minimum number of nuclei per perturbation cutoff for downstream analyses. **(D)** Scatterplot showing single gRNA assignment correlation to discrepancy for FACS labeling rate between left and right hemispheres within one animal (discrepancy in FACS gating between samples). **(E)** Histogram of 8,588 gRNA distribution in AAV library and scatter plot of gRNA correlation between two different batches of viral preparation. **(F)** Ranked bar plot of total number of nuclei assigned with single perturbation identity for each binned cell type. **(G)** Ranked bar plot of total number of nuclei assigned with single perturbation identity for each binned cell type, colored by perturbation identity.

**Figure S5.**
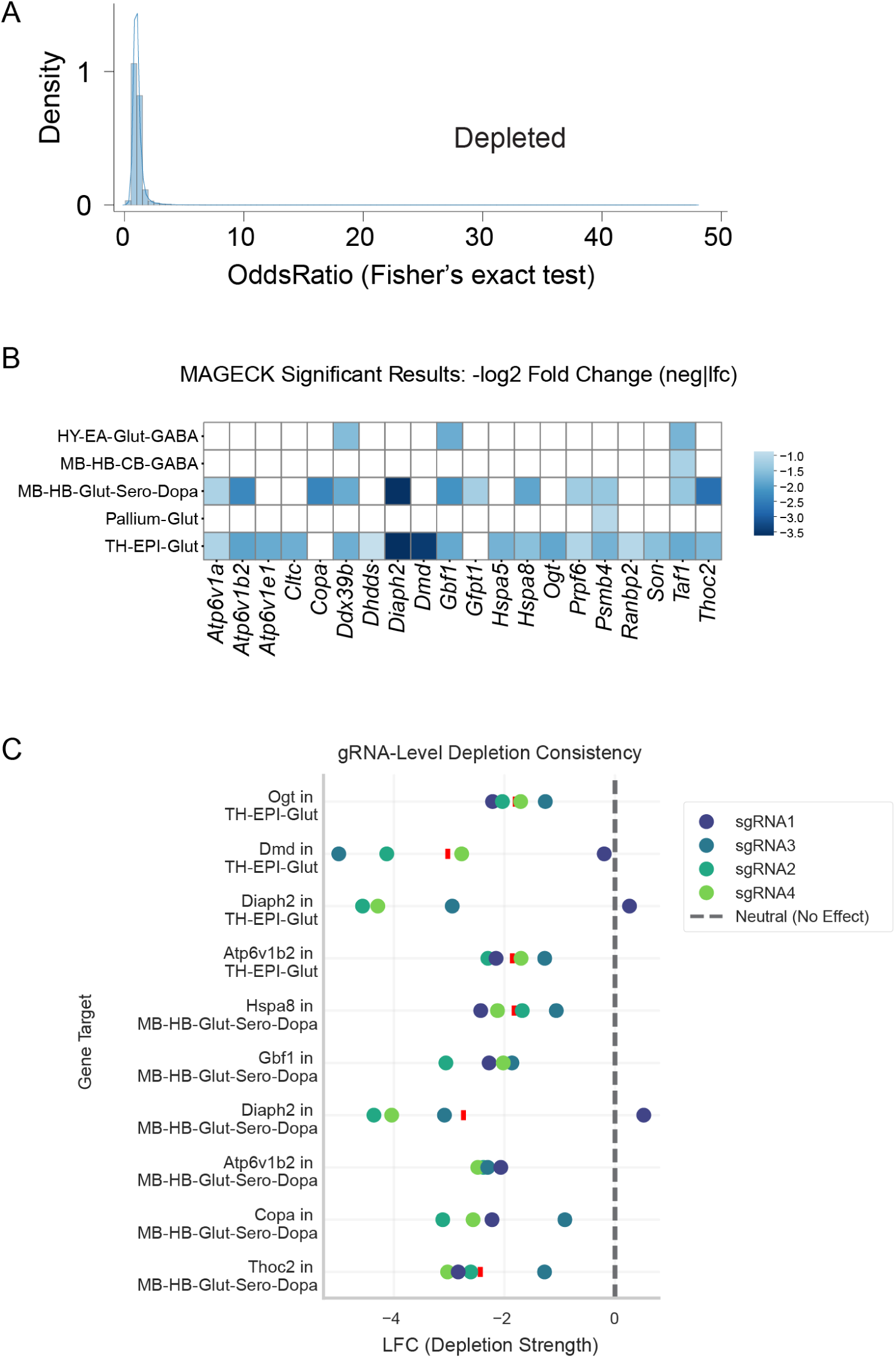
Cell type depletion analysis with Fisher’s exact test and MAGeCK. **(A)** Density plot of odds ratios for all depleted perturbation–cell type pairs using Fisher’s exact test. **(B-C)** MAGeCK significant results (log₂ fold-change) for depleted perturbation–cell type pairs across binned neuronal classes, concordant with Fisher’s exact test results shown in Fig. 2A-C.

**Figure S6.**
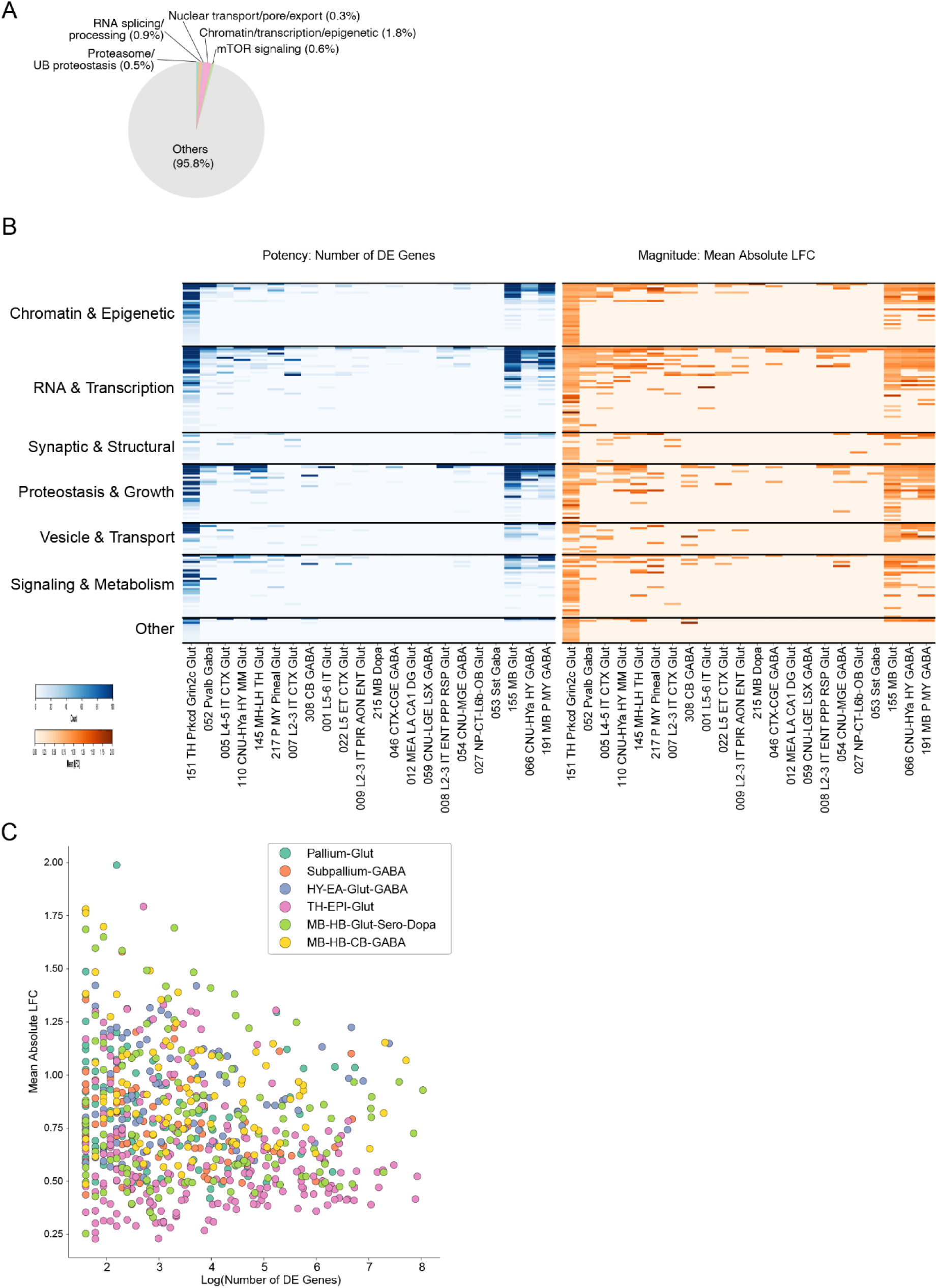
Pathway composition and perturbation effect structure across cell types. **(A)** Pie chart showing pathway category composition among all 1,947 perturbed genes, with the vast majority (95.8%) falling outside the core pathway categories that dominate among top-DEG perturbations (compared to Fig. 2D), indicating that specific molecular systems are disproportionately represented among high-impact perturbations. **(B)** Heatmaps showing perturbation potency (number of DE genes; left, blue) and magnitude (mean absolute log-fold-change; right, orange) for each perturbation across cell types, organized by functional pathway category. Each row represents a single perturbation; columns represent binned neuronal cell types. **(C)** Scatter plot of perturbation potency (log number of DE genes) versus magnitude (mean absolute LFC) across all perturbation–cell type pairs, colored by binned neuronal class. Points in the upper-left quadrant reflect perturbations with few but large-magnitude DEGs (cell-type-restricted, strong effects), while points in the lower-right reflect perturbations with many but modest-magnitude DEGs (broad, distributed effects).

**Figure S7.**
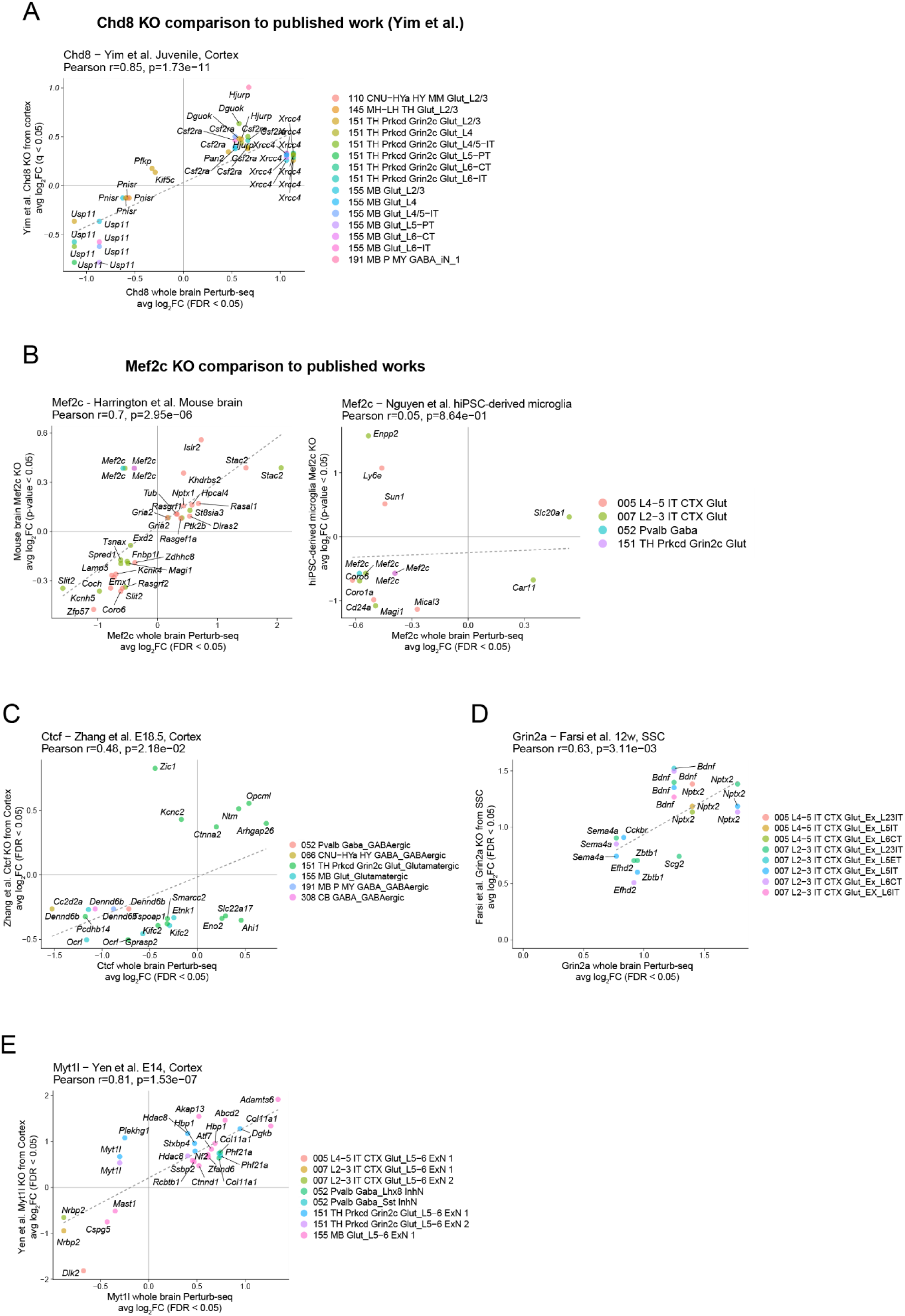
in vivo Perturb-seq DEG alignment with published germline or conditional knockout data. **(A)** *Chd8* knockout comparison with a reference *Chd8* KO dataset^16^ (Yim et al., juvenile cortex). Log fold changes (logFC) of DEGs (FDR < 0.05) from the whole-brain perturb-seq dataset are compared against logFC from the reference for overlapping DEGs (q-value < 0.05). **(B)** *Mef2C* knockout comparisons with published datasets. Left: Harrington et al.^17^ (mouse brain). Right: Nguyen et al.^21^ (hiPSC-derived microglia). LogFC from the whole-brain perturb-seq dataset are compared against logFC from references for overlapping DEGs (p-value < 0.05). **(C-E)** Knockout comparison with reference datasets for *Ctcf* ^18^ (Zhang et al., E18.5 cortex) **(C)**, *Grin2a*^19^ (Farsi et al. 12-week somatosensory cortex) **(D)**, *Myt1l*^20^ (Yen et al., E14 cortex) **(E)**. LogFC from the whole-brain perturb-seq dataset are compared against logFC from each reference for overlapping DEGs (FDR < 0.05).

**Figure S8.**
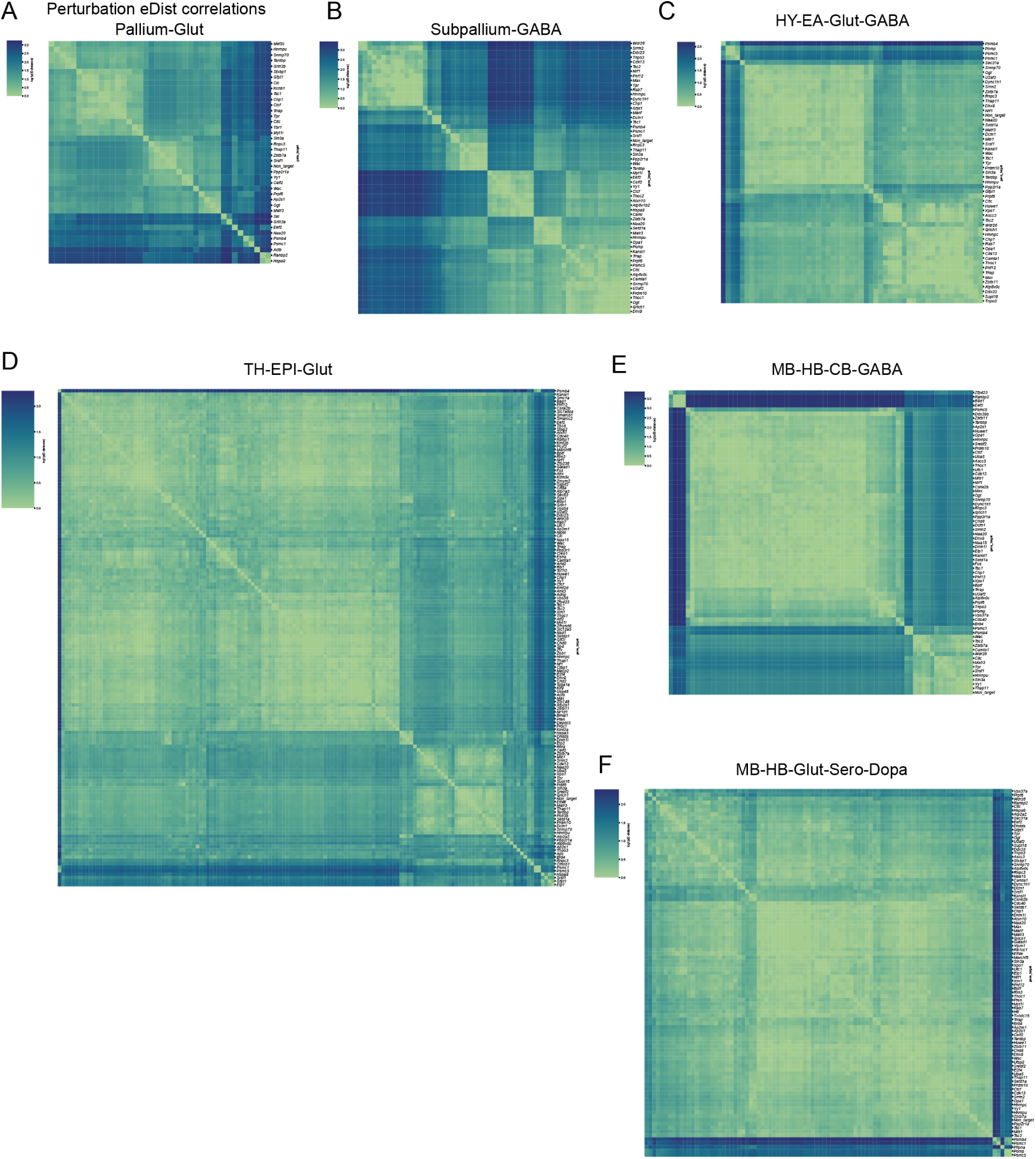
Cell-type-specific perturbation energy distance matrices. **(A–F)** Pairwise energy distance correlation matrices for effective perturbations computed within individual binned neuronal classes: Pallium-Glut (A), Subpallium-GABA (B), HY-EA-Glut-GABA (C), TH-EPI-Glut (D), MB-HB-CB-GABA (E), and MB-HB-Glut-Sero-Dopa (F). Perturbations are hierarchically clustered within each cell type. While modular structure is apparent across all classes, the specific clustering patterns differ between neuronal types, indicating that perturbation relationships are shaped by cell-type context. These cell-type-specific patterns motivated further investigation of perturbation similarity at the level of individual perturbation–cell type pairs (Fig. 3B).

**Figure S9.**
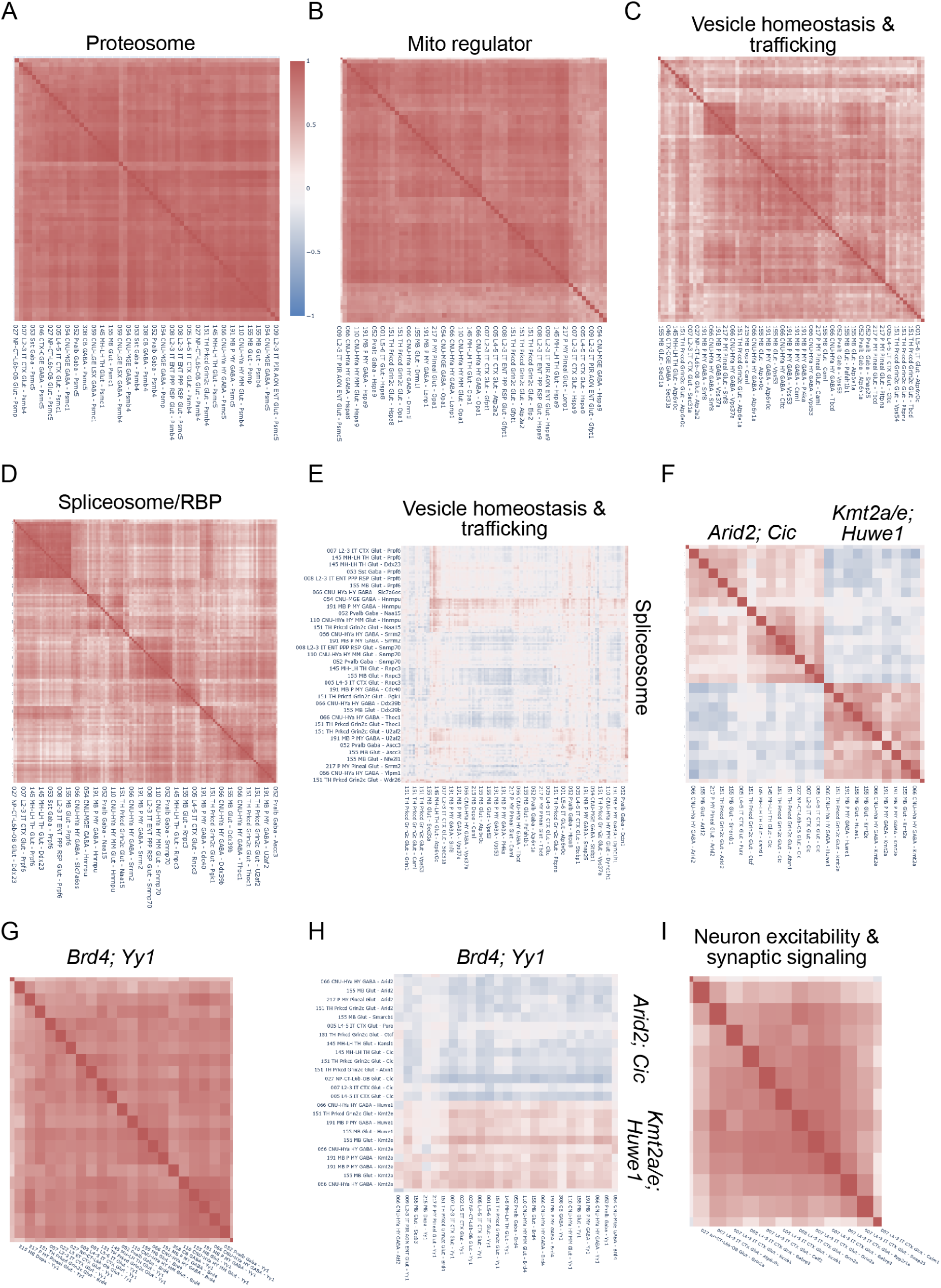
Cosine similarity matrices for perturbation modules identified by hierarchical clustering. **(A)** Pairwise cosine similarity among proteasome-related perturbation–cell type pairs, showing high internal concordance. **(B)** Mitochondrial quality-control regulators (*Hspa5*, *Hspa8*, *Hspa9*, *Lonp1*, *Opa1*, *Dnm1l*), forming a tightly correlated module. **(C)** Vesicle homeostasis and trafficking module, including endosomal and secretory machinery components (*Vps37a*, *Vps53*, *Snf8*, *Atp6v0c*, *Atp6v1a*, *Sec31a*, *Pitpna*, *Tbcd*). **(D)** Spliceosome and RNA-binding protein module (*Prpf6*, *Snrnp70*, *Rnpc3*, *U2af2*, *Srrm2*). **(E)** Cross-module similarity between vesicle homeostasis/trafficking and spliceosome modules, showing antagonistic (negative) correlation between these functional classes. Rows and columns are labeled by perturbation–cell type pair. **(F)** Chromatin remodeling sub-modules, including SWI/SNF components and histone modifiers (*Kmt2a/e*, *Huwe1*) and a separate *Arid2*/*Cic* cluster, showing related but distinct transcriptional signatures. **(G)** *Brd4*/*Yy1*-associated transcriptional activation module. (H) Cross-module similarity between *Brd4*/*Yy1* and chromatin repressor modules (*Arid2*/*Cic*, *Kmt2a/e*/*Huwe1*), revealing an antagonistic axis between transcriptional activation and repressive chromatin regulators. **(I)** Neuronal excitability and synaptic signaling module (*Grin2a*, *Kcnb1*, *Gabrg2*, *Snap25*, *Celf2*, *Ppp2r1a*), with cross-module relationships to chromatin sub-modules shown.

**Figure S10.**
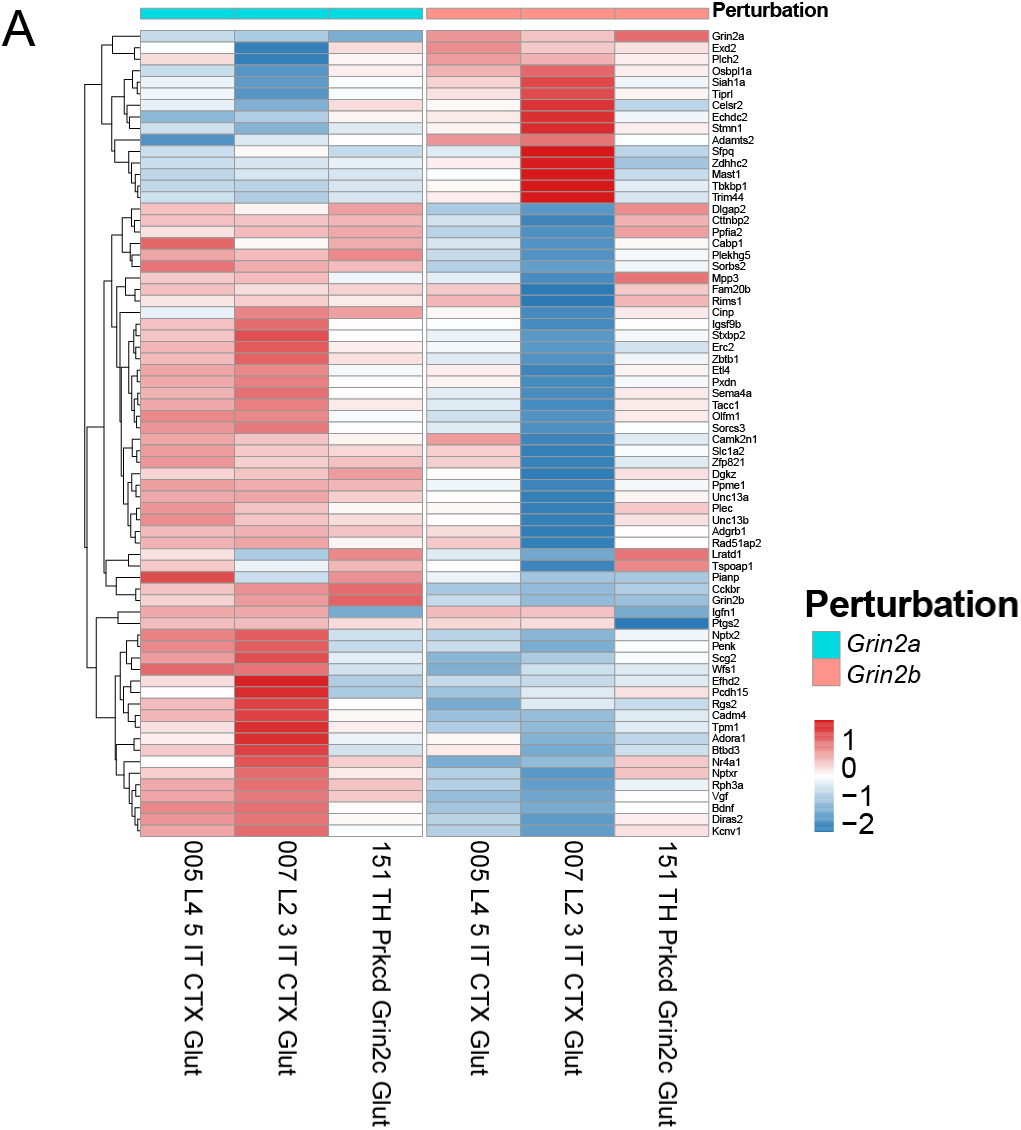
Context-specific perturbation effects of Grin2a and Grin2b. **(A)** Heatmap of selected DEGs showing variable opposing regulation between Grin2a and Grin2b perturbations in different cell types.

**Figure S11.**
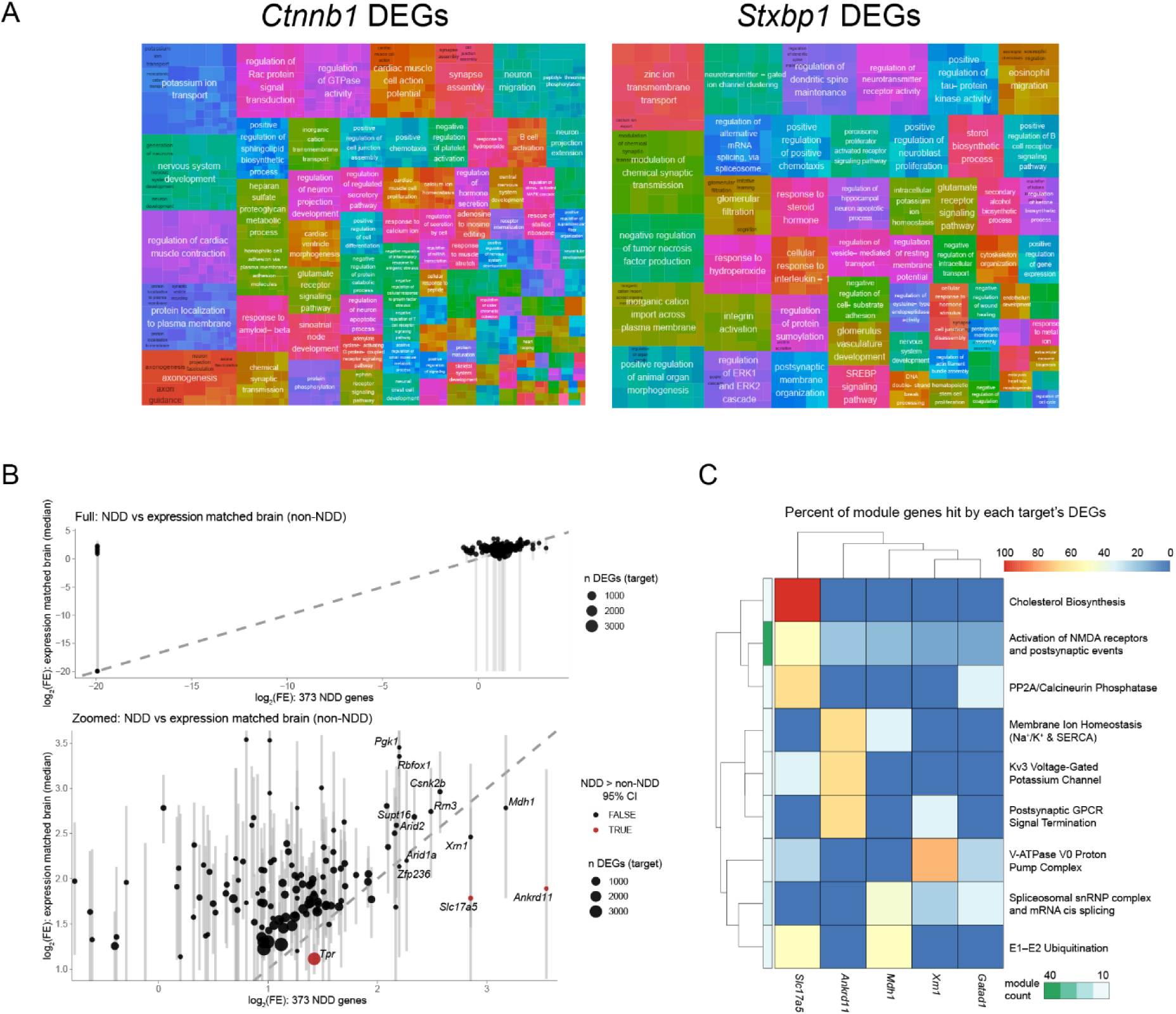
NDD gene enrichment and functional convergence of perturbation targets. **(A)** Treemap visualization of Gene Ontology Biological Process terms enriched with DEGs under perturbation of Ctnnb1 (left) and Stxbp1 (right). **(B)** Enrichment of downstream DEGs with NDD risk genes (n = 373) compared to an expression-matched brain background. Each point represents a perturbation target; point size reflects the number of DEGs. The dashed line indicates equivalence between enrichment against NDD genes and matched controls. The lower panel shows a zoomed view highlighting individual targets. Error bars represent 95% empirical confidence intervals (CI) from 1,000 bootstraps. Targets with full 95% CIs for enrichment of expression-matched genes smaller than enrichment for NDD genes are in red. **(C)** Heatmap showing percent of module genes per cluster identified in the STRING PPI network.

## Notes

### Competing Interest Statement

MK, SR, SD, TJC, and LS are employees of NVIDIA. DD and GXYZ are employees of PerturbAI. GXYZ and XJ are co-founders of PerturbAI. SJS receives research funding from BioMarin Pharmaceutical.

### Funding

- National Institute of Health: R01MH125516, R01HG012819, R01MH137042
- Medical Research Council Centre of Research Excellence in Therapeutic Genomics: MR/Z504725/1
- Health Data Research UK QQ2 Molecules to Health Records Driver Programme
- Simons Foundation: SFARI #736613
- California Institute for Regenerative Medicine: DISC4-16295
- Howard Hughes Medical Institute
- G. Harold & Leila Y. Mathers Foundation
- One Mind Foundation
- Conrad Prebys Foundation
- Pew Charitable Trusts
- McKnight Foundation
- James Fickel and Jed McCaleb

